# A selective alternative pathway complement inhibitor for treatment of paroxysmal nocturnal hemoglobinuria

**DOI:** 10.1101/2024.06.23.600249

**Authors:** Hagen Sülzen, Hanna Tulmin, Miroslav Hájek, Jaroslav Čermák, Alžběta Kadlecová, Petr Pompach, Jitka Votrubová, Martin Zoltner, Sebastian Zoll

## Abstract

The complement system, a critical component of the human innate immune system, enhances the ability to clear microbes and damaged cells. Dysregulation of this system, particularly the alternative pathway (AP), can lead to several rare blood disorders such as paroxysmal nocturnal hemoglobinuria (PNH). This study introduces SH-01, a blood parasite-derived, novel recombinant protein which selectively inhibits the AP. We show that SH-01 effectively prevents the lysis of erythrocytes isolated from PNH patients. Unlike current treatments such as eculizumab, SH-01 targets the AP without impairing the classical or lectin pathways, reducing the risk of infections and extravascular hemolysis. SH-01 functions through a unique two-stage mechanism, preventing C3b deposition and inhibiting AP C5 convertase activity while maintaining the amplification loop’s functionality. Immunization studies in mice showed no significant immune response against SH-01, and the protein exhibited high stability and no acute toxicity. These findings suggest SH-01 as a promising candidate for treatment of PNH and other diseases characterized by AP hyperactivation, offering a more targeted therapeutic and thus safer approach.

## INTRODUCTION

The complement system, a highly efficient proteolytic activation cascade of plasma proteins, is an integral and vital component of the human innate immune system, enhancing the ability of antibodies and phagocytes to swiftly clear microbes, damaged cells, and promote inflammation. However, if not adequately regulated, due to genetic abnormalities or external factors, the complement system’s ability to distinguish between host and foreign materials may be compromised. This can lead to a variety of severe diseases, many of which have a genetic background and can be fatal is left untreated(*1*). Briefly summarized, the complement cascade is initiated through three distinct pathways: the classical pathway (CP), the lectin pathway (LP), and the alternative pathway (AP). All three pathways converge in the formation of membrane-bound C3 convertases(*2*) (Figure 1).

**Figure 1.**
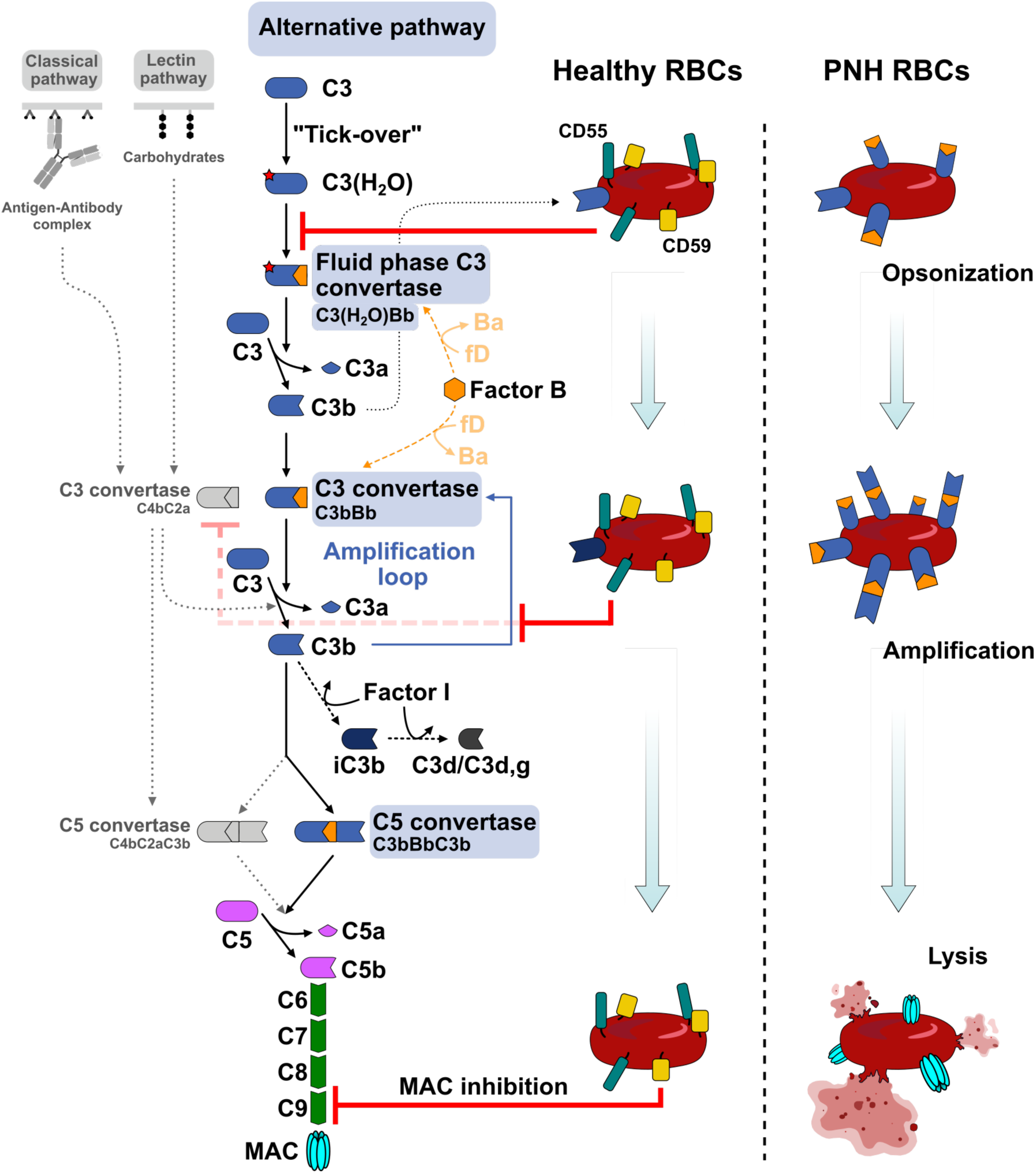
Overview of the alternative complement pathway and its negative regulators. Contrary to the antigen-driven CP and LP, the AP is initiated spontaneously by hydrolysis of C3 (“tick-over”), subsequently forming the AP fluid phase C3 convertase. Newly generated C3b is deposited on pathogen surfaces where it nucleates formation of the AP C3 convertase, amplifying the complement activity. Binding of an additional C3b molecule to the C3 convertase forms the C5 convertase, initiating C5 cleavage and thereby progression to the terminal pathway, culminating in the formation of the MAC. On healthy erythrocytes, CD55 inhibits formation and decreases stability of the C3 convertase of AP and CP/LP. For simplicity, the inhibition of the CP/LP C3 convertase is only shown illustratively and a visualization of accelerated decay of the convertases has been omitted. CD59 exerts its inhibitory function by preventing incorporation of MAC components C8 and C9. On PNH erythrocytes devoid of CD55, C3b is readily deposited, and the complement activity is amplified, resulting in progression to the terminal pathway. A lack of CD59 fails to prevent formation of the MAC, resulting in hemolysis. Abbreviations: RBC - red blood cell, Ba - factor B fragment a, Bb - factor B fragment b, fD – factor D, MAC – membrane attack complex.

While the CP and LP are antigen-driven, triggered by a range of pathogen surface moieties, the AP, in addition to amplifying complement activity after CP/LP activation (via the so-called “amplification loop”), is continuously activated at a low level due to the spontaneous hydrolysis of the mildly unstable complement factor C3 (“tick-over”)(*3*). This activity results in formation of the fluid-phase C3 convertase which cleaves native C3 to C3b, the central hub of all 3 pathways(*4*). In the AP, C3b binds to cell surfaces where it nucleates the formation of the AP C3 and C5 convertases (C3bBb and C3bBbC3b), initiating the terminal pathway (TP), which leads to assembly of the membrane attack complex (MAC) and, ultimately, cell lysis (Figure 1).

During progression of the CP and LP, C4b substitutes for C3b, forming the CP/LP C3 convertase with C2a (C4bC2a). Subsequent binding of an additional C3b molecule results in the CP/LP C5 convertase (C4bC2C3b). Despite their different origins and structures, both convertase types cleave C3 and C5(*2*) (Figure 1).

Since the AP is activated spontaneously in absence of foreign microbial surfaces and conjugation of C3b to hydroxyl groups on cellular surfaces is too unspecific to be controlled directly, the complement activity must be tightly regulated by additional factors to protect host cells from its potentially harmful effects. In healthy individuals, regulation is accomplished by a variety of host-derived modulators that prevent the assembly of surface-bound C3b into functional convertases and thereby ultimately inhibit the formation of the MAC. On resting cells, negative regulation of the AP occurs immediately after the deposition of C3b through the action of factor I (fI) and its cofactors factor H (fH) and CD35 (Complement Receptor 1, CR1), which convert C3b into iC3b and eventually C3d,g. These proteolytic derivatives are no longer able to nucleate the formation of convertases (*2, 5–8*)(Figure 1).

On activated cells, C3 convertases have already formed and must be disassembled to prevent further progression of the cascade. A multitude of surface-bound negative regulators, such as CD35, CD46 (Membrane Cofactor Protein, MCP), CD55 (Decay Accelerating Factor, DAF) and CD59 (MAC-inhibitory protein, MAC-IP) play crucial roles in this process(*2, 9–12*).

Regardless of the complement system’s importance for averting infections, its dysregulation, particularly that of the AP, has been linked to a series of severe diseases, including, but not limited to Alzheimer’s disease, autoimmune conditions, schizophrenia, nephropathies, macular degeneration, and Crohn’s disease(*13*). Prototypical examples of complement dysregulation disorders include atypical hemolytic uremic syndrome (aHUS), C3 glomerulopathy (C3G), and paroxysmal nocturnal hemoglobinuria (PNH)(*1*).

The predominant causes for aHUS are mutations in the thioester domain (TED) of C3b, factor B (component of the AP C3 convertase) and across various negative regulators of the AP, such as fH, fI, and CD46(*14–17*). These mutations lead to a loss of cellular self-recognition and uncontrolled lysis of erythrocytes, ultimately resulting in kidney failure.

C3G, on the contrary, is caused by mutations that affect fluid-phase regulation of the AP and lead to complement deposition within the glomerulus(*18*).

PNH is associated with loss-of-function mutations in glycosyltransferase genes required for synthesizing glycosylphosphatidylinositol (GPI), resulting in a deficiency of GPI-anchored membrane proteins such as the complement inhibitors CD55 and CD59(*19–21*). CD55, also commonly known as ‘decay-accelerating factor’ (DAF), inhibits formation and decreases the stability of both, the AP and CP/LP C3 convertases (Figure 1). Contrastingly, CD59 specifically targets the TP, the final stage of complement activation, effectively blocking membrane perforation by inhibiting the assembly of MAC components C5b-8 and C5b-9 (Figure 1). Both proteins are crucial for the regulation of complement activity and their deficiency results in hemolysis (Figure 1), causing severe pathologies such as thromboembolism and chronic kidney disease(*22*).

Due to the unambiguous and well-established role of complement dysregulation in the pathophysiology of PNH, the pathological severity and the relative ease with which response to a treatment can be quantified, PNH has become the paragon of diseases considered for testing new complement inhibitors(*1, 23, 24*).

The current gold-standard for treatment of PNH and similar indications, such as aHUS, is eculizumab (Soliris, Alexion Pharmaceuticals), a therapeutic monoclonal antibody that targets complement component C5 and inhibits the activation of the TP. Eculizumab effectively prevents intravascular hemolysis of erythrocytes in patients with PNH, leading to a rapid improvement in their conditions.

However, by acting downstream of the convergence point for the CP, LP, and AP, all three complement pathways are inhibited to the same extent, increasing the risk of contracting bacterial infections, particularly for *Neisseria* spp., some strains of which can cause meningitis. Since eculizumab does not target the early steps of complement activation, opsonization with inactivated C3b derivates (C3d,g and iC3b) still occurs, thereby potentially leaving patients susceptible to extravascular hemolysis(*25, 26*).

Furthermore, the efficacy of eculizumab treatment may vary considerably due to the diverse underlying mechanisms that result in disease manifestation, as exemplified by the drug’s limited success in treatment of C3G(*27*).

In C3G, the primary driver of the disease is excessive generation and deposition of C3 breakdown products in the renal glomerulus, leading to kidney damage. The root cause of C3G therefore lies with complement factors upstream of C5, explaining the limited efficacy of a TP inhibitor in treatment of this condition.

Here we introduce SH-01, a novel, parasite-derived recombinant protein that selectively inhibits the alternative complement pathway through a distinct mechanism of action and is capable of efficiently protecting PNH erythrocytes from complement-mediated lysis.

Previously, we have elucidated the structure of the parent molecule of SH-01, ISG65, for which we proposed an initial mechanism of action, demonstrating that its interaction with C3b can fully inhibit AP-mediated hemolysis whereas CP and LP function remains unimpaired(*28*).

In this study, we report the extended mechanism of action for SH-01. We demonstrate that its inhibition of the AP occurs through a unique two-stage mechanism that involves direct prevention of C3b deposition on the cell surface and inhibition of the activity of AP C5 convertases, all while retaining functionality of the amplification loop after CP/LP activation.

Immunization of mice with SH-01 elicited no significant immune response against the C3 binding site. Instead, it induced a focused antibody response to a hotspot of functionally irrelevant epitopes which are likely to serve as immune decoys on the parasite surface. Furthermore, SH-01 demonstrated high stability in human plasma and exhibited no acute toxicity in mice, rendering it a promising drug candidate for treatment of PNH and various other diseases characterized by hyperactivation of the alternative pathway. Without compromising the unaffected pathways of the innate immune response, SH-01’s unparalleled specificity could contribute significantly to patient safety during treatment.

## MATERIAL AND METHODS

### C3b deposition ELISA

C3b deposition was assessed using the Wieslab complement system alternative pathway kit (Svar Life Science, Sweden). Non-heat inactivated normal human serum (Sigma-Aldrich) was diluted 1:18 (100%) and 1:36 (50%) using the kits included dilution buffer. 1.5 µL SH-01 (2.8 mg/mL) was added to the serum samples at a final concentration of 1 µM. For control samples, an equivalent volume of dilution buffer was added. Following the addition of SH-01/dilution buffer, serum samples were incubated for 15 min at RT, while shaking. After transferring the samples to the Wieslab ELISA plate, the plate was incubated at 37°C for 60 min, as per manufacturer instructions. Following the removal of the samples after the incubation step, the wells were washed four times with PBS (10 mM phosphate buffer, 140 mM NaCl, 3 mM KCl, pH 7.4) containing 0.05% Tween-20 (=PBST). Deposition of C3b was subsequently detected by incubation with a rabbit polyclonal anti-C3c-FITC IgG (1:100 dilution in PBST) (ab4212, Abcam) for 1 hour at RT, protected from light. After the incubation, samples were removed, and the ELISA plate washed as described previously. Finally, 100µL PBST were added to the wells and fluorescence was analyzed using an Infinite M1000 microplate reader (Tecan) with excitation and emission wavelengths of 495 nm and 519 nm, respectively. All samples were measured in technical triplicates. C3b deposition was illustrated with GraphPad Prism 10 (GraphPad Software).

### C3b deposition analysis by flow cytometry

C3b deposition on IgG sensibilized sheep erythrocytes (CH50 assay, HaemoScan) was assessed via flow cytometry. Sheep erythrocytes were washed according to manufacturer’s instructions. Complement C6 depleted human serum (C1288, Sigma-Aldrich), diluted to final concentration of 2% with dilution buffer (CH50 assay, HaemoScan), was incubated with SH-01 at final concentrations ranging from 0 to 50 µM. The samples were incubated for 25 min at RT before adding them to 25 µL of erythrocyte suspension (containing ∼2.5 × 10^6^ cells) to a total volume of 50 µL. Erythrocytes were incubated in the depleted serum at 37°C for 15 min. The incubation was stopped by addition of 500 µL of ice-cold dilution buffer and samples were centrifuged for 5 min, 400x g, 4°C. Erythrocyte pellets were resuspended in 100 μL ice-cold dilution buffer supplemented with 0.2% bovine serum albumin (BSA, Sigma-Aldrich) and stained on ice for 1 hour with a monoclonal anti-iC3b antibody (A209, Quidel), conjugated with AF_647_ (ab269823, Abcam) (2.4 ng/µL final concentration), and polyclonal anti-C3c-FITC IgG (ab4212, Abcam) (1:300 final dilution) while protected from light. Flow cytometry was performed on a CytoFLEX LX flow cytometer (Beckman Coulter), data analysis was conducted with the CytExpert software (Beckman Coulter). Fractions of positive singlets were illustrated with GraphPad Prism 10 (GraphPad Software).

### Flow cytometry-based competition assay

#### Preparation of fluorescently labelled antigen and antigen-complexes

SH-01 was dialyzed into 0.2M sodium bicarbonate buffer, 150 mM NaCl, pH 8.3 and subsequently concentrated to 8 mg/mL. AF_594_ NHS ester (Invitrogen) was resuspended to 10 mg/mL in anhydrous DMSO and added to SH-01 to a final concentration of 1 mg/mL. After 1h at RT, the reaction was quenched by addition of tris(hydroxymethyl)aminomethane to a final concentration of 200 mM. Unreacted dye was removed by size exclusion chromatography (SEC) using Superdex 200 Increase 10/300 GL column (GE Healthcare), pre-equilibrated in HBS (20 mM HEPES, 150 mM NaCl, pH 7.5). Fractions containing SH-01_AF594_ were collected and pooled. The complexes of SH-01_AF594_ with C3d and Fab 3E12 were obtained by concentrating ∼1 mg of either protein with 300µg of SH-01_AF594_ to ∼200µL before removing the excess protein via SEC on a Superdex 200 Increase 10/300 GL column (GE Healthcare), pre-equilibrated in HBS. Fractions containing both SH-01_AF594_ and C3d/Fab were identified by SDS-PAGE analysis, pooled and concentrated to ∼1 mg/mL before flash-freezing in liquid nitrogen. Samples were subsequently stored at -80°C until use.

#### Isolation of B-cells

Spleens from mice immunized with 1200 µg SH-01 / 30g body weight (BW) were harvested on day 12 after the immunization, fatty tissue removed and the isolated spleens stored on ice.

1.5 h after sacrifice of the animals, B-cells were isolated using a gentleMACS tissue dissociator (Miltenyi Biotec) in combination with the mouse spleen dissociation kit and C-tubes (Miltenyi Biotec), according to the manufacturer’s instructions. Erythrocytes were lysed using ACK lysis buffer (10 mM potassium bicarbonate, 155 mM ammonium chloride, 0.1 mM EDTA, pH 7.4). The remaining cells were washed and resuspended in PBS for staining with Zombie NIR dye (BioLegend). After staining with the viability marker, the cells were washed and resuspended in ‘staining buffer’ (PBS, 1% fetal bovine serum (v/v)). FcR Blocker (Cat. No. 130-092-575, Miltenyi Biotec) was added according to the manufacturer’s instructions. After washing and resuspending the cells in staining buffer, anti-CD19-FITC (BioLegend) (1:10 (v/v) and SH-01_AF594_ (19 µg/mL final) were added and incubated for 30 mins at RT, protected from light. After the incubation period, cells were washed and resuspended in staining buffer. Positively labelled, vital B-cells singlets (Zombie NIR -, CD19-FITC +, SH-01_AF594_ +) were gated for and sorted into RPMI-16 media using a CytoFLEX cytometer (Beckmann Coulter), supplemented with 1% (v/v) low-endotoxin FBS. Final cell concentration was approximately 2 × 10^6^ cells/mL. Cells were cultured in sterile T-flasks at 37°C ON.

#### Flow cytometry analysis of isolated B-cells

Isolated B-cells were harvested by centrifugation (350x g, 10 min, RT) and washed three times with PBS. After the final washing step, cells were re-suspended in 100 µL PBS. Cells were split into 25 µL aliquots and 1 µL of PBS, SH-01_AF594_, SH-01_AF594_:C3d or SH-01_AF594_:Fab were added (∼40 µg/mL final).

The samples were subsequently analyzed for AF_594_ positive cells, using a CytoFLEX cytometer with the CytExpert software (v 2.3.1.22, Beckman Coulter). Sufficient washing was confirmed by absence of the anti-CD19-FITC signal (data not shown).

### Generation of mAb 3E12 and Fab 3E12

#### Mouse immunizations

Female BALB/c mice were injected with 200 μg of antigen mixture intraperitoneally in a total volume of 200 μL (i.e., 100 μL of antigen at 2 mg/mL with 100 μL complete Freund’s adjuvant (CFA, Sigma)). It was crucial to prepare a stable emulsion to generate a strong immune response, which was achieved by vortexing the mixture for several minutes before drawing it into a syringe. After 21 days, the mice received a booster injection with the same amount of recombinant protein, but CFA was substituted with IFA (incomplete Freund’s Adjuvant, Sigma). After 42 days, the BALB/c mice were boosted intravenously via the tail vein with 5 μg of antigen suspended in 100 μL PBS. After 45 days, the mice were sacrificed, and their serum and spleens were harvested.

#### Generation of Hybridoma cell lines

Splenocytes were extracted from a single spleen and fused with Sp2/0 myeloma cells. The resulting hybridoma cells were selected on semi-solid methylcellulose and grown in 96-well plates using the ClonaCell-HY Hybridoma Cloning Kit (Stemcell Technologies). Hybridomas secreting SH-01-reactive IgG were identified by several rounds of ELISA screening. Hybridoma lines expressing monoclonal antibodies were cultured in Dulbecco’s Modified Eagle Medium (DMEM, Sigma-Aldrich) supplemented with 4 mM L-glutamine (Sigma-Aldrich), 100 U/mL penicillin, 0.1 mg/mL streptomycin (Sigma-Aldrich), and 10% fetal calf serum (Gibco). The cells were then gradually transferred to serum-free CD Hybridoma medium (Life Technologies) and cultured in suspension. After 14 days, the cells were harvested, and the supernatant containing the secreted IgG was ultra-filtered and dialyzed against 20 mM HEPES (pH 8.0).

#### Purification of monoclonal antibodies and Fab generation

ELISA-positive monoclonal IgG was further tested for binding to SH-01 by analytical size exclusion chromatography. To purify monoclonal antibodies, 1 L of the hybridoma supernatant was first concentrated using tangential flow filtration (TFF) and eluted from the cassette with Buffer A (20 mM Tris, pH 7.5). Protein A/G agarose was equilibrated with 50 mL of Buffer A. The buffer-exchanged hybridoma supernatant was then applied to the equilibrated slurry twice. The agarose beads were washed with 300 ml of Buffer B (20 mM Tris, 300 mM NaCl, pH 8.0) and eluted with 20 mL of Buffer C (0.1 M glycine, pH 2.5). Fractions of 2 ml were collected and immediately neutralized by adding 120 µL of Buffer D (1 M Tris, pH 9.0), ensuring the final pH was in the neutral range.

For Ficin cleavage of mouse IgG1 to generate Fabs, the purified mAb was dialyzed against 5 L of Buffer E1 (100 mM citrate, pH 6.0) and concentrated to 5 mg/mL. A 50 ml solution of 10x Digestion Buffer F1 (100 mM citrate, 50 mM EDTA, 250 mM cysteine) was prepared and adjusted to pH 6.0. Ficin Agarose (2 mL of settled beads) was activated by washing with 2x20 mL of 1x Digestion Buffer F1 and centrifuged for 1 minute at 1000 × g. The IgG1 in 100 mM citrate buffer was mixed with 10% of 10x Digestion Buffer F1 and added to the activated Ficin Agarose. The mixture was incubated at 37°C for 3-5 hours while shaking at high speed. If cleavage was incomplete after the maximum time, the incubation was continued overnight. The Ficin Agarose was separated from the IgG fragments using a gravity flow column, washed with an equal volume of Buffer G (10 mM Tris, pH 7.5), and combined with the digest.

#### Peptide microarray

Peptide microarrays and experiment were manufactured and performed by JPT Peptide Technologies GmbH (Berlin, Germany). Briefly summarized, a library of 15-residue-long peptides with a 13-residue overlap covering the entire sequence of SH-01, was synthesized and immobilized on microarray slides (139 peptides total). Full-length human and mouse IgG were co-immobilized on the microarray slides as assay controls.

Purified mouse mAb 3E12 was diluted to a final concentration of 1 µg/mL with Blocking Buffer (SuperBlock T20 (TBS) Blocking Buffer, Thermo Fisher) and incubated with the microarrays, previously washed with washing buffer (50 mM TBS, 0.1% Tween-20, pH 7.2, JPT Peptide Technologies GmbH), for 1 hour at 30°C. Following sample incubation, the microarrays were washed with washing buffer and incubated with DyLight650-labeled anti-mouse IgG (Abcam ab97018) (1 µg/mL final) for 1 hour at 30°C. Control incubations with only secondary antibody (no primary antibody) were performed to assess non-specific binding. After incubation with the secondary antibody, the microarrays were washed with washing buffer and dried before scanning.

The microarrays were scanned using an automated microarray scanner system (GenePix SL50 slide loader & GenePix 4300, Molecular Devices) operated at 635 nm. The resulting images were analyzed and quantified using spot-recognition software GenePix (Molecular Devices). For each peptide, the mean signal intensity was extracted (between 0 and 65535 arbitrary units). Subsequently, MMC2 values were determined to ensure consistent data quality. If the coefficient of variation, which is calculated by dividing the standard deviation by the mean, is greater than 0.5, the MMC2 is equal to the mean of each of the three microarray instances. In this instance, the MMC2 is assigned the mean of the two closest values (MC2). Finally, background fluorescence induced by unspecific binding of the secondary antibody in absence of a primary antibody was subtracted. Fluorescence intensity was illustrated using GraphPad Prism 10 (GraphPad Software).

#### SAXS data collection and processing

Small-angle X-ray scattering (SAXS) data were collected at the BioSAXS beamline P12 at DESY in Hamburg, Germany, using a Pilatus 6M detector (DECTRIS) with a wavelength of 1.24 Å at 20 °C. For SEC-SAXS measurements, 50 µL of SH-01 in complex with Fab 3E12 at a concentration of 7.2 mg/ml were injected onto a Superdex 200 5/150 GL column. The column was pre-equilibrated with 20 mM HEPES, 150 mM NaCl, and 3% (v/v) glycerol (pH 7.5), and the flow rate set to 0.3 mL/min. Scattering data were collected as the components eluted from the column and passed through the SAXS measuring cell. The ATSAS software package was utilized to normalize the data to the intensity of the incident beam, average the frames, and subtract the scattering contribution from the buffer. Specifically, 10 frames corresponding to the void volume of the column were averaged and subtracted from 10 averaged frames of the main elution peak. The radius of gyration (Rg), maximum particle dimension (Dmax), and distance distribution function (p(r)) were evaluated using the program PRIMUS, part of the ATSAS package(*29*).

### Hydrogen-deuterium exchange mass spectrometry

Hydrogen-deuterium exchange was initiated by 10-fold dilution of the SH-01 or SH-01:Fab3E12 in a deuterated buffer (20 mM Hepes, 150 mM NaCl, pD 7.5). After 20 s incubation in the deuterated buffer, 50 microliters of quench buffer (3 M Urea, 1 M Thiourea, 1 M glycine, 200 mM TCEP at pH 2.3) was added and immediately injected on a Nepenthesin-2/pepsin column (AffiPro, Czech Republic) by an automated sample handling system. Generated peptides were desalted by a trap column (Luna Omega 5 um Polar C18 100 Å Micro Trap 20 × 0.3 mm) for 3 min at a flow rate 200 µL min^-1^ using isocratic pump delivering 0.4% formic acid in water. After 3 min, peptides were separated on a C18 reversed phase column (Luna® Omega 1.6 μm Polar C18 100 Å, 100 × 1.0 mm) and analyzed using a timsToF Pro mass spectrometer. Both protease column and trap column were placed at 4°C. Peptides were separated by linear gradient 10-30% B in 18 min. The mass spectrometer was operated in positive MS mode. Spectra of partially deuterated peptides were processed by Data Analysis 5.3 (Bruker Daltonics, Billerica, MA) and by in-house program DeutEx(*30*).

### Analytical size exclusion chromatography

100 µg of SH-01 was incubated with a three-fold molar excess of Fab 3E12 (350 µg), concentrated to ∼50 µL using a Vivaspin 2 spin-concentrator with 10kDa MWCO (Cytiva), and subjected to SEC on a Superdex 200 3.2/300 Increase column (Cytiva), pre-equilibrated with running buffer (20 mM HEPES, 150 mM NaCl, pH 7.5) and connected to an Äkta Pure Micro FPLC system (Cytiva).

300 µL fractions were collected throughout the elution. Peak fractions were analyzed by reducing SDS-PAGE analysis. The elution fractions containing the desired SH-01:Fab 3E12 complex were pooled and subsequently incubated with an a three-fold molar excess of C3d (100 µg), concentrated to ∼50 µL subjected to SEC again, as described before. Peak elution fractions were once more analyzed by reducing SDS-PAGE.

### Acute toxicity of SH-01

The acute toxicity study of SH-01 after intraperitoneal (i.p.) administration in mice was conducted at the Institute of Physiology, Department of Biological controls of the Czech Academy of Sciences (OBK IPHYS CAS) in Prague (Czech Republic). The testing facility is accredited under the Certificate of Good Laboratory Practice for Medicinal Products for Human Use (Act No. 378/2007 Coll.), and although this study was not conducted in GLP mode, it was performed in accordance with established Quality Assurance Program standards. All procedures adhered to the Standard Operating Procedures (SOPs) of the OBK IPHYS CAS, consistent with the European Convention for the Protection of Vertebrate Animals

Used for Experimental and Other Scientific Purposes (ETS 123) and the Czech Collection of Laws No. 246/1992, including subsequent amendments. The facility’s operation and animal care protocols are further validated by the Accreditation Certificate MZE-23481/2021-18134, issued by the Ministry of Agriculture of the Czech Republic. Ethical approval for the study was granted by the Institutional Animal Care and Use Committee (IACUC) and the Committee for Animal Protection of the Czech Academy of Sciences (Approval No. 55/2019), ensuring compliance with the Animal Welfare Act and adherence to OECD and 3R principles.

The study was performed on 15 male C57BL/6 JRccHsd mice (Envigo – AnLab s.r.o.). The animals, 8 weeks old upon arrival at the test facility, were divided into four experimental dose groups and a control group which received a single i.p. administration of the vehicle (PBS). Each study group contained 3 animals. The mice were acclimatized for 7 days before the start of the study, only animals in good health were used. The first dose of SH-01 (400 µg / 30g BW) was administered in a single i.p. injection in a total volume of 500 µL. An equal volume of the vehicle (PBS, pH 7.4) was administered in identical fashion to the control group. In absence of clinical signs of toxicity after an initial 72-hour observation period, subsequent dose groups received progressively higher doses of 800 µg, 1200 µg, and 2000 µg of SH-01 per 30g BW, with each increase preceded by another 72-hour evaluation period to ensure that no signs of acute toxicity occurred.

All mice were frequently observed for clinical signs (such as signs of toxicity, changes in skin, fur, eyes and mucous membranes; respiratory-, circulatory-, autonomic- and central nervous system; somatomotor activity and behavioral patterns) during the first 30 minutes after dosing, then once more after 1 h, 2 h, 3 h and 24 hours after dosing.

Subsequently, clinical signs were assessed twice daily for a total of 12 days. All animals were weighed individually before randomization, before administration of SH-01 as well as on day 4, day 8 and day 12 after administration. All animals were euthanized on day 12 and a gross necropsy performed.

### Plasma stability of SH-01

#### Time course experiment

SH-01 (18.9 mg/mL) was added to 500 µL of normal human plasma to a final concentration of 2 mg/mL. After thorough mixing, the aliquot of the reference sample (Day 0) was taken immediately and diluted into PBST (1:20), before flash freezing in liquid nitrogen. All frozen aliquots were stored at -80°C until further use.

The experiment-tube, closed and sealed with parafilm, was constantly incubated at 37°C throughout the remainder of the experiment. Before removing sample at the given time points, the sample was vortexed gently and briefly centrifuged (<500x g) to collect any condensate from the lid. The appropriate sample volume was removed and diluted into PBST (1:20) before flash freezing in liquid nitrogen. Samples for all following timepoints were acquired as described.

#### Western blot

The frozen aliquots were thawed simultaneously in a heated water bath at 37°C before any precipitate was removed by centrifugation (21.000x g, 4°C, 5 min). 12 µL of the cleared supernatant were combined with 3 µL of 6x non-reducing loading dye before boiling (100°C, 1 min). The samples were centrifuged (21.000x g, RT, 3 min) before all 15 µL of each sample were loaded onto a 12.5% polyacrylamide gel for SDS-PAGE (43 min, RT, 220 V constant). Proteins were transferred onto a Nitrocellulose membrane, pre-soaked in transfer buffer (20 mM Tris, 190 mM glycine, 20% Methanol, pH 8.3) using a Mini-Transfer cell (BioRad) for 1.5 hours (100 V constant, 4°C).The membrane was transferred into SuperBlock T20 (TBS) blocking buffer (Thermo Scientific) and blocked overnight while shaking at 4°C. After removing the blocking solution, the membrane was rinsed and washed three times for 5 minutes with PBST. The membrane was subsequently incubated with polyclonal anti-SH-01 mouse serum (*see Generation of mAb 3E12 and Fab fragment*) (1:100 final, 2 h, RT). After removing the polyclonal serum, the membrane was rinsed and washed three times as described previously before incubation with the IRDye 680RD conjugated donkey anti-mouse IgG secondary antibody (LI-COR Biosciences) (1:10.000 final, 1.5h, RT, dark). From this point onwards, the membrane was protected from light. After the incubation period, the membrane was washed three times, as described, before imaging.

#### Analysis/Quantification

The Western Blot was imaged wet on an Odyssey CLx (LI-COR Biosciences) with the ImageStudio Software (Version 5.2.5). Fluorescence was detected using the 700 nm channel. Fluorescence intensity was analyzed with the ImageJ software. In brief, an area encompassing the entire reference band (Day 0) was selected and brightness measured within. The same area-size and shape was used to measure the brightness of the other bands. For each lane, the background signal was determined above the measured protein band using the same area. The final intensity was obtained by subtracting the measured brightness of the background from the respective protein band. The percentage of fluorescence intensity was calculated relative to the reference sample.

### Isolation of PNH erythrocytes and ABO-matched human serum

#### Human blood samples

Peripheral blood samples were obtained from two patients (patients A and B) receiving regular maintenance doses of eculizumab (900 mg Soliris (AstraZeneca), diluted in 120 mL saline solution, administered once every 2 weeks) for treatment of PNH, and from a single, untreated patient diagnosed with PNH (patient C). PNH patient blood was collected in EDTA vacutainer tubes (ref. no. 367525, BD). Blood samples from the treated patients were obtained immediately before administration of the next scheduled maintenance dose.

Blood samples from healthy volunteers to obtain ABO-matched normal human serum were collected in vacutainer tubes for serum analysis (ref. no. 367896, BD).

Venipuncture was performed according to standard operating procedures of the Institute of Hematology and Blood Transfusion (Ústav hematologie a krevní transfuze) (Prague, Czech Republic) following informed consent from voluntary donors, as approved by the local Institutional Review Board and in accordance with the Declaration of Helsinki.

#### Isolation of PNH RBCs and normal human serum

Shortly after the blood-draw, PNH patient blood samples were centrifuged (450x g, 10 min, 4°C) and the plasma and buffy coat removed. Plasma of patients undergoing eculizumab treatment was aliquoted and flash-frozen in liquid nitrogen. The pelleted erythrocytes were resuspended and subsequently washed 3 times in ice cold SAGM (saline-adenine-glucose-mannitol) solution. Washed erythrocytes were stored in SAGM at ∼ 50% hematocrit for maximally 5 weeks at 4°C. Following the centrifugation of blood samples from healthy individuals (1350x g, 15 min, 4°C), the serum was removed, aliquoted and immediately flash-frozen in liquid nitrogen. Serum samples were stored at -80°C until further use.

### Flow cytometry analysis of PNH erythrocytes

Preceding incubation with PNH phenotype erythrocytes, normal human serum (NHS) was initially pre-incubated with SH-01 (48 µM final concentration, 25 min, RT). Alternative complement pathway activation was initiated by acidification, as described previously(*31*). Briefly summarized, the SH-01:NHS solution was supplemented with MgCl_2_ (1.5 mM final) and acidified (aNHS) to pH 7 by addition of 300 mM HCl (1:16.7 (v/v) final). 5 × 10^6^ PNH patient erythrocytes were pelleted by centrifugation (450x g, 5 min, 4°C), the supernatant removed and the cell pellet gently resuspended in 50 µL of acidified SH-01:NHS solution. Following the incubation period (37°C, 1 hour), RBCs were washed twice with ice cold RBC staining buffer (PBS supplemented with 0.2% BSA) before staining on ice with anti-CD59_AF405_ monoclonal antibody (FAB1987V, R&D Systems, 5 ng/µL final concentration) and anti-C3c-FITC polyclonal antibody (ab4212, Abcam, 1:300 final) for 1 h.

After the staining, unbound antibodies were removed by washing the erythrocytes with RBC staining buffer twice. Hemolysis and C3b deposition were subsequently assessed by flow cytometry analysis using a CytoFLEX cytometer with the CytExpert software (Beckman Coulter). Baseline levels of CD59-negative cells and C3b deposition were established by incubating erythrocytes in RBC staining buffer instead of aNHS. RBCs incubated in ABO-matched aNHS served as a positive control for hemolysis and efficient opsonization. RBCs incubated in acidified plasma from PNH patients undergoing eculizumab treatment served as a positive control for efficient inhibition of hemolysis of PNH phenotype erythrocytes.

## RESULTS

### SH-01 blocks C3b deposition in the AP, but does not affect the CP/LP, and prevents degradation to iC3b

We previously showed that SH-01 fully inhibits hemolysis via the AP but has no inhibitory effect on complement-mediated lysis via the CP/LP(*28*). We could demonstrate that SH-01 diminishes the activity of assembled AP C5 convertases by binding to C3b but noted that for complete inhibition, as observed in hemolysis assays with rabbit erythrocytes, an additional interference mechanism must be present.

Using a solid phase immunoassay for AP activity, we show here that the addition of 1 µM of SH-01 to normal human serum (NHS) fully prevents C3b surface deposition through binding of C3b in the fluid phase (Figure 2A). This discovery, in conjunction with the inhibition of AP C5 convertase, provides an explanation for the previously observed two-stage mechanism by which SH-01 achieves complete inhibition of hemolysis via the AP.

**Figure 2.**
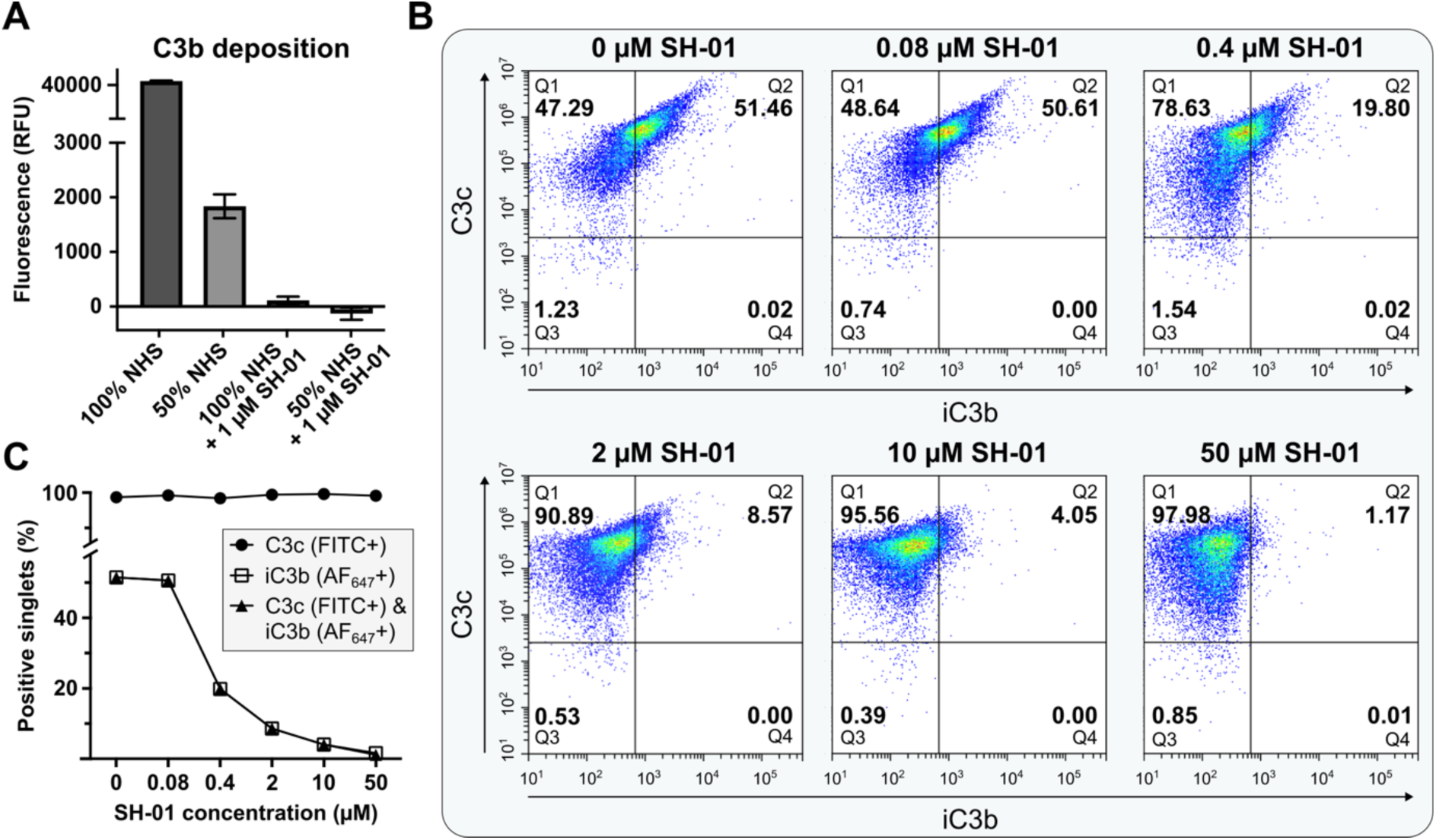
SH-01 does not inhibit C3b deposition during CP/LP activation but prevents degradation to iC3b. **A** Solid phase immunoassay measuring the deposition of C3b through activation of the alternative complement pathway (n = 2 technical replicates; data points are shown as mean values ± SD). Addition of 1 µM SH-01 to NHS completely abrogates C3b deposition. **B** Flow cytometry analysis of C3b deposition via activation of the classical complement pathway. Bivariate density plots, divided into four quadrants, illustrate presence of C3b, iC3b and C3d,g on single sheep erythrocytes (Q1: C3b^+^; Q2: iC3b^+^; Q3: C3b^-^, iC3b^-^ & C3d,g ^-^; Q4: C3d,g^+^). Percentage of RBCs in each quadrant is indicated. >98 % of analyzed cells were positive for C3c across all concentrations of SH-01 added. The addition of 50 µM SH-01 decreased the number of iC3b positive cells in a concentration dependent manner, reducing the fraction of positive from 51.46 to 1.17 %. **C** Summary of flow cytometry analysis (shown in panel B).

Next, we dissected the impact of SH-01 on the CP/LP. Since C3b is not a component of the CP/LP C3 convertase C4bC2a, SH-01 was expected to exhibit no effect on deposition or activity of this convertase. Our previous findings corroborated this assumption, demonstrating that SH-01 does not prevent CP/LP mediated hemolysis(*28*).

However, analogue to the AP C3 convertase, C3 does also serve as the substrate for C4bC2a, which subsequently converts it to C3b, the primary target of SH-01. We therefore investigated whether SH-01 could also prevent the deposition of membrane proximal C3b, generated by the cell surface-associated CP/LP C3 convertase. By performing flow cytometry analysis with sensitized sheep erythrocytes incubated in C6-depleted human serum, we could demonstrate that SH-01 does indeed not prevent C3b deposition in this setting, even at concentrations up to 50 µM (Figure 2B,C). C3b deposition was visualized using a polyclonal antibody against the integral C3c domain (Supplementary Figure 1A).

In absence of SH-01, deposited C3b is readily converted to iC3b, as demonstrated by detection with a conformation-specific anti-iC3b antibody (Figure 2B,C, Supplementary Figure 1B). It should be noted that while the used antibody does not recognize C3b, it fails to differentiate between iC3b and its proteolytic cleavage product C3d,g. However, the lack of C3c in the C3d,g fragment allows for unambiguous identification of all three fragments when staining for both, iC3b and C3c.

Addition of SH-01 to the C6-depleted serum prior to incubation with the erythrocytes revealed a concentration-dependent inhibition of the conversion of C3b to iC3b (Figure 2B,C, Supplementary Figure 1B). By inhibiting the proteolytic degradation to iC3b, deposited C3b could allow for formation of AP C3 convertases, thereby maintaining functionality of the amplification loop following activation of CP/LP, even in presence of SH-01 (Figure 1). Consequently, this effect would also effectively prevent the opsonization of cells via iC3b. While further research is needed to elucidate the precise mechanism by which SH-01 inhibits iC3b conversion, structural alignments indicate that SH-01 blocks the binding site for the CCP19 domain of factor H (fH) on C3b (Supplementary Figure 2A). Previous research has shown that binding of CCP19 to the TED is crucial for the correct alignment of fH domains CCP1-4 on C3b(*32, 33*). These domains act as a cofactor for factor I (fI)-mediated conversion of C3b to iC3b on the cell surface (Supplementary Figure 2B). It therefore seems likely SH-01 driven abrogation of fH binding by SH-01 contributes to the observed prevention of iC3b generation.

### SH-01 does not evoke a neutralizing antibody response against the C3b binding site

When used as therapeutic agents in human patients, biologics of non-human origin, such as SH-01, are likely to provoke an immune response. Depending on their immunogenicity, the adaptive immune system may raise drug-specific antibodies. These antibodies can either facilitate the removal of the drug from the bloodstream through phagocytic uptake or neutralize its therapeutic effects by binding to and obstructing its active site(s). To obtain a better understanding of the immunological characteristics of SH-01, we set out to identify particularly immunogenic epitopes.

In order to do so, spleens from mice immunized with SH-01 were harvested and B-cells isolated. These antibody-presenting cells were then analyzed in a flow cytometry-based competitive binding assay, utilizing AlexaFluor-594 labeled SH-01 (SH-01_AF594_) alone, SH-01_AF594_ in complex with C3d (the proteolytically liberated TED of C3b), and SH-01_AF594_ in complex with a mAb-derived Fab (3E12) from a hybridoma cell line established in a previous immunization cycle (Figure 3).

**Figure 3.**
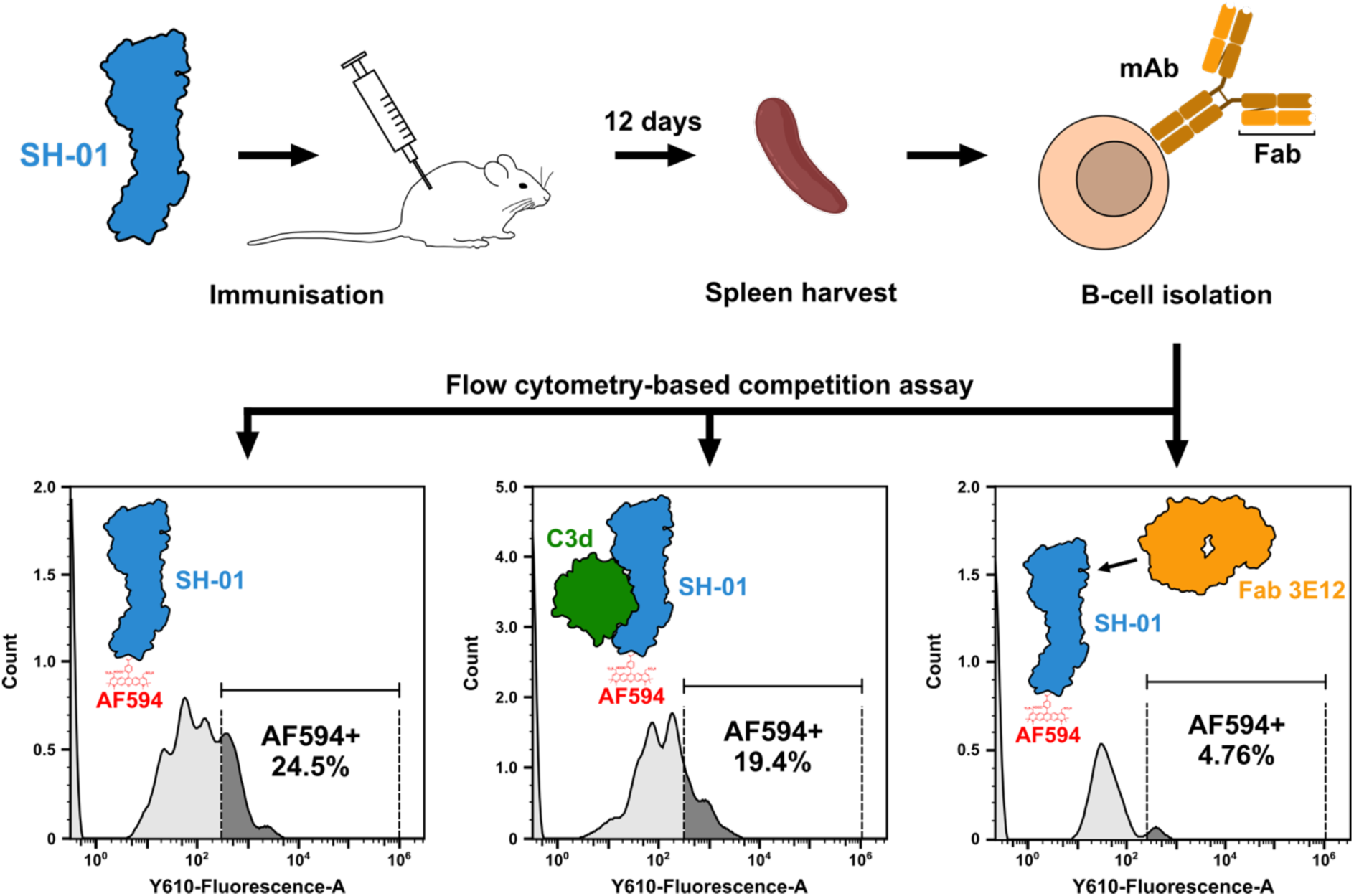
Flow cytometry-based competition assay to characterize the immunological profile of SH-01. Spleens from mice immunized with SH-01 were harvested 12 days after challenge with the antigen. Subsequently isolated B-cells were incubated with SH-01_AF594_, SH-01_AF594_:C3d or SH-01_AF594_:3E12, washed and analyzed for AF594 using flow cytometry. 24.5% of cells incubated with SH-01_AF594_, were positive for AF594. Whereas Incubation with SH-01_AF594_:C3d resulted only in a marginal decrease in positive cells, incubation with SH-01_AF594_:3E12 caused a significant decrease of positive cells to 4.76%.

Incubation of antibody presenting B-cells with SH-01_AF594_ resulted in 24.5% of AF594-positive cells. Saturation of the C3b binding site of labelled SH-01 with C3d did not significantly change the number of AF594 positive cells (19.4%) in comparison to the cells incubated with SH-01 alone. This indicates that most presented antibodies were non-neutralizing for the interaction with C3b. However, when SH-01_AF594_ was saturated with Fab 3E12, only 20% of the initial B-cell population size was detected as AF594 positive (4.8%) (Figure 3) (Supplementary Figure 3).

This finding suggests that binding by the Fab could either shield antibody binding to other immunogenic epitopes on SH-01 or induce structural changes that expose neo-epitopes, which in turn are not recognized by the antibodies presented on the B-cells.

### The head domain of SH-01 is an antibody binding site and immune-dominant decoy

Monoclonal antibody 3E12 was the only high-affinity binder identified through a hybridoma selection process. This, in conjunction with its ability to reduce the binding of other antibodies to the antigen, suggested that it may recognize a potentially unique epitope on SH-01. To identify this epitope, we mapped the binding of Fab 3E12 to SH-01 using a peptide microarray and determined the envelope structure of the Fab-bound complex through small-angle X-ray scattering (SAXS) (Figure 4, Supplementary Table 1).

**Figure 4.**
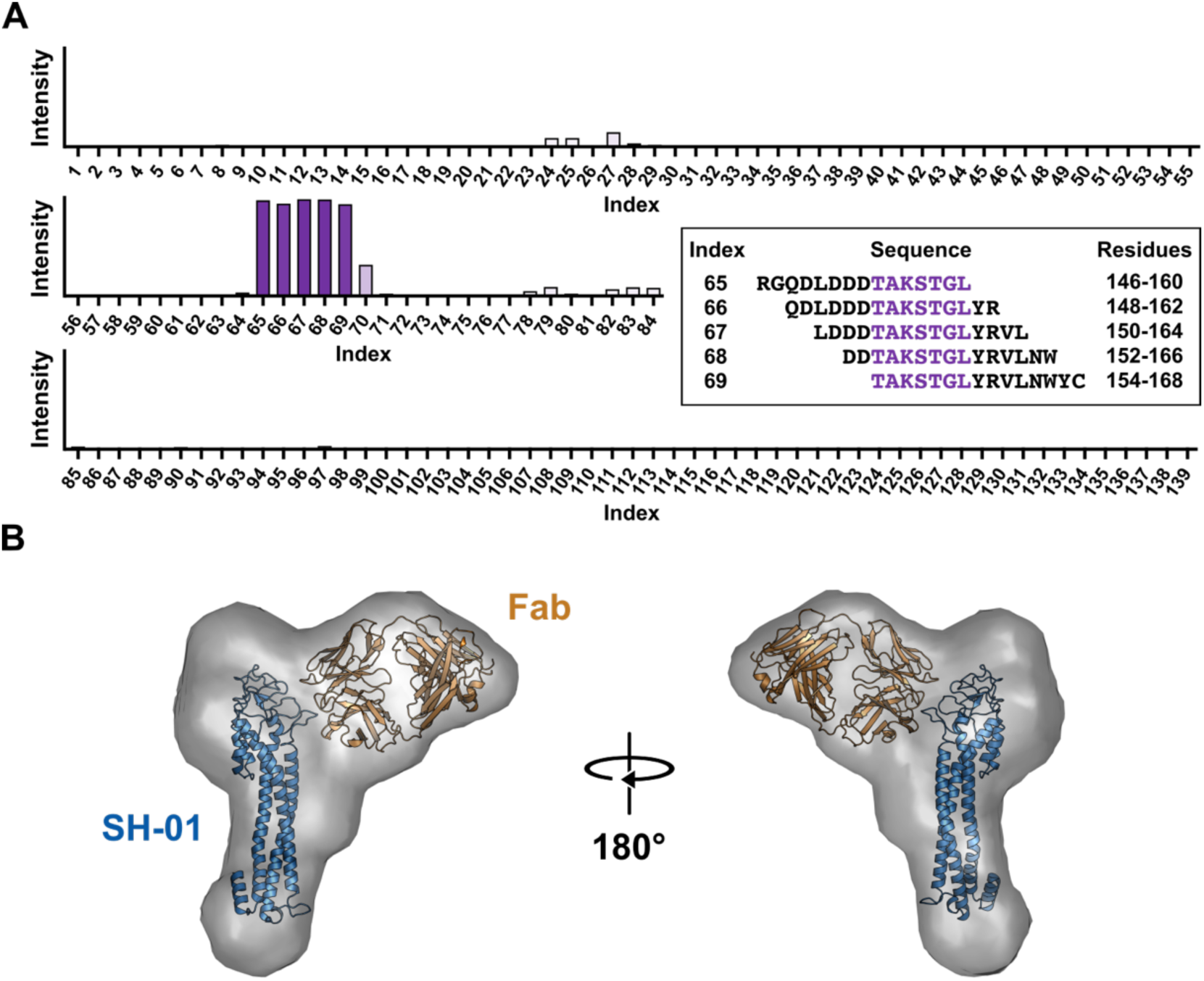
Fab 3E12 binds an epitope in the SH-01 head domain. **A** Peptide microarray for mapping the Fab 3E12 epitope on SH-01. The bar diagram shows mean fluorescence intensity for each peptide. Peptide sequences with the highest signal intensity are shown in the inset, the likely consensus sequence, TAKSTGL, is highlighted in purple. Residue numbers indicate position of the corresponding peptide in the sequence of SH-01. **B** SAXS envelope of the SH-01:Fab 3E12 complex, confirming binding of the Fab to SH-01 in vicinity of the TAKSTGL sequence. SH-01 is shown as blue cartoon. Since no experimental structure for 3E12 is available, a representative model of a mouse-derived Fab, shown in orange, has been used instead (PDB 3VG0).

For the peptide array, a series of 15-residue-long peptides with a 13-residue overlap, spanning the entire sequence of SH-01, was immobilized on a glass surface via a hydrophilic linker moiety. Detection of Fab 3E12 binding with a fluorescently labelled anti-mouse IgG antibody distinctly identified a stretch of seven residues (TAKSTGL), locating the Fab epitope within the loop-rich head domain of SH-01 (Figure 4A). In line with this finding, the SAXS envelope confirmed the Fab’s binding in this region. Given the distinctive conformations of both proteins, the Fab could be confidently docked into the space adjacent to the head domain of SH-01, where it effectively shields a large surface area. These results demonstrate that epitopes within the head domain of SH-01 are viable antibody binding sites.

### Binding of Fab 3E12 introduces conformational changes in SH-01

Interaction of a biologic with a patient’s antibody repertoire may directly impact the drug’s therapeutic efficacy. Consequently, we sought to elucidate the effects of 3E12 binding on the structure of SH-01 and investigated whether antibody binding to the head domain exerts an allosteric effect on its interaction with C3. Hydrogen-deuterium exchange mass spectrometry (HDX-MS) appeared to be the most suitable method for observing any potential structural changes in SH-01. HDX-MS has no size limitations and does not require labor-intensive and time-consuming structure determination. By measuring the relative exchange-rate of amide hydrogens to deuterium in a D_2_O-containing buffer, information about the relative solvent exposure of individual protein regions can be obtained(*34*).

Solvent exposure could change in response to either transitions between structured and unstructured regions or due to ligand binding. Deuteration typically occurs at a faster rate in unstructured regions than in well-structured counterparts and is reduced in areas where ligand binding shields deuterium incorporation. By using HDX-MS, we could confirm binding of Fab 3E12 to SH-01 in the TAKSTGL sequence that we initially identified using a peptide microarray (Figure 5A, Supplementary Figure 4).

**Figure 5.**
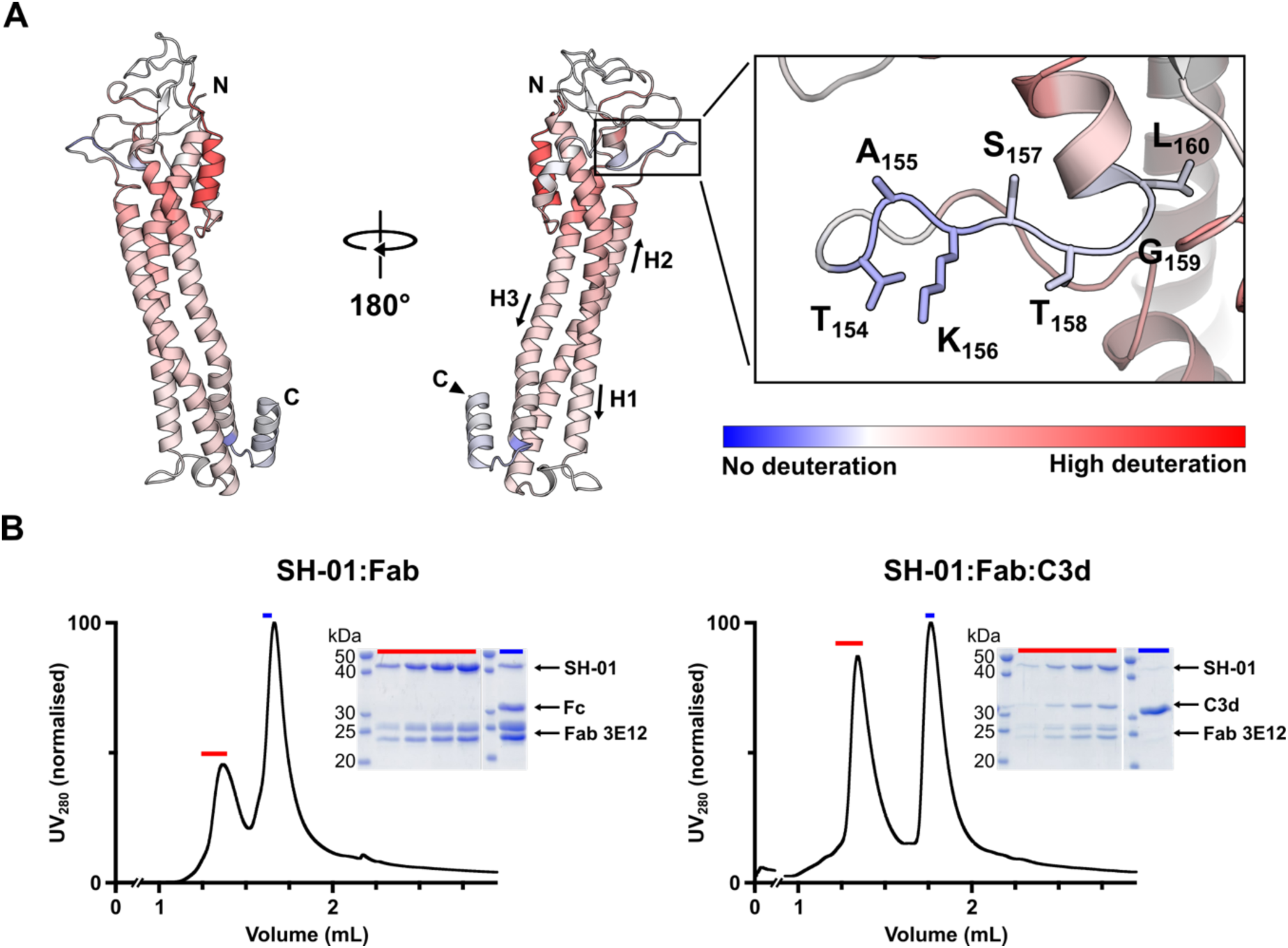
Antibody binding to SH-01 induces structural changes but does not impair binding to its therapeutic target. **A** Deuteration of SH-01 in presence of Fab3E12, as determined by HDX-MS, mapped onto a model of SH-01. The color gradient indicates the relative deuteration level after 20s of incubation in deuteration buffer. Areas with reduced deuteration, such as the Fab 3E12 epitope, shown in detail in the inset, are colored in blue. Areas of high deuteration are colored in red. N- and C-terminus of the SH-01 model are indicated. The disordered C-terminus of the SH-01 model has been removed for clarity. The helices constituting the three-helical bundle of SH-01 are numerated and their directionality indicated with arrows. **B** Saturation of SH-01 with Fab3E12 does not preclude binding to C3d (i.e. the thioester domain of C3b) in size exclusion chromatography as demonstrated by SDS-PAGE analysis of peak fractions, highlighted in red and blue in the chromatogram. The chromatograms have been normalized to the maximum observed absorbance at 280 nm for clarity.

Binding of the Fab to SH-01, effectively shielding the binding interface from the solvent, resulted in a clear reduction in deuterium uptake of the linear epitope when compared to the unbound structure.

Interestingly, the epitope is located at a “handle”-shaped loop insertion, breaking up helix 2 (H2) of the 3 main helices that constitute the three helical bundle of SH-01, thereby separating a smaller helix from its longer stem and displacing it to the head domain (Figure 5A).

While a reduction in deuteration rates was limited to the precise Fab 3E12 binding site on SH-01, surprisingly, other regions of the protein exhibited a dramatically increased deuterium uptake upon binding (Figure 5A). This increase was particularly pronounced towards the head domain of SH-01 which we have previously demonstrated to be prone for antibody binding (Figure 3). While HDX-MS does not allow to infer information on the exact structure of the SH-01:3E12 complex, it clearly demonstrated an allosteric effect of Fab 3E12 binding on the overall protein conformation. In conjunction, these findings indicate that SH-01 adopts a noticeably less well-structured conformation in complex with the Fab, incorporating deuterium more rapidly than its unbound counterpart.

To investigate whether this allosteric effect may impair SH-01’s ability to bind its therapeutic target, C3b, we performed analytical size exclusion chromatography (SEC). First, SH-01 was incubated with a three-fold molar excess of 3E12, subjected to SEC, and analyzed by reducing SDS-PAGE. The elution fractions corresponding to the complex were pooled, incubated with a three-fold molar excess of C3d and subjected to SEC again. Peak elution fractions were once more analyzed by reducing SDS-PAGE. Coomassie staining revealed four bands corresponding to SH-01, C3d, and the heavy and light chains of Fab 3E12 (Figure 5B, Supplementary Figure 5). This demonstrated that Fab-bound SH-01, despite the large-scale conformational changes induced by the binding of 3E12, can still interact with the thioester domain of C3b with high affinity.

### Plasma stability and acute toxicity of SH-01

High plasma stability and tolerability (and conversely lack of toxicity) are essential qualities for a therapeutic agent. Here, we show that SH-01 meets both of these criteria. In an *ex vivo* stability assay, SH-01 was incubated with human plasma at physiological temperature and sampled daily over an 11-day period. The presence of SH-01 was then assessed by Western blotting using polyclonal anti-SH-01 mouse serum and a fluorescently labeled secondary anti-mouse IgG antibody. Protein amounts were determined by band densitometry relative to day 0. SH-01 exhibited high stability in human plasma, with 63% of the intact full-length protein remaining on day 11, demonstrating little susceptibility to proteolytic cleavage (Figure 6A).

**Figure 6.**
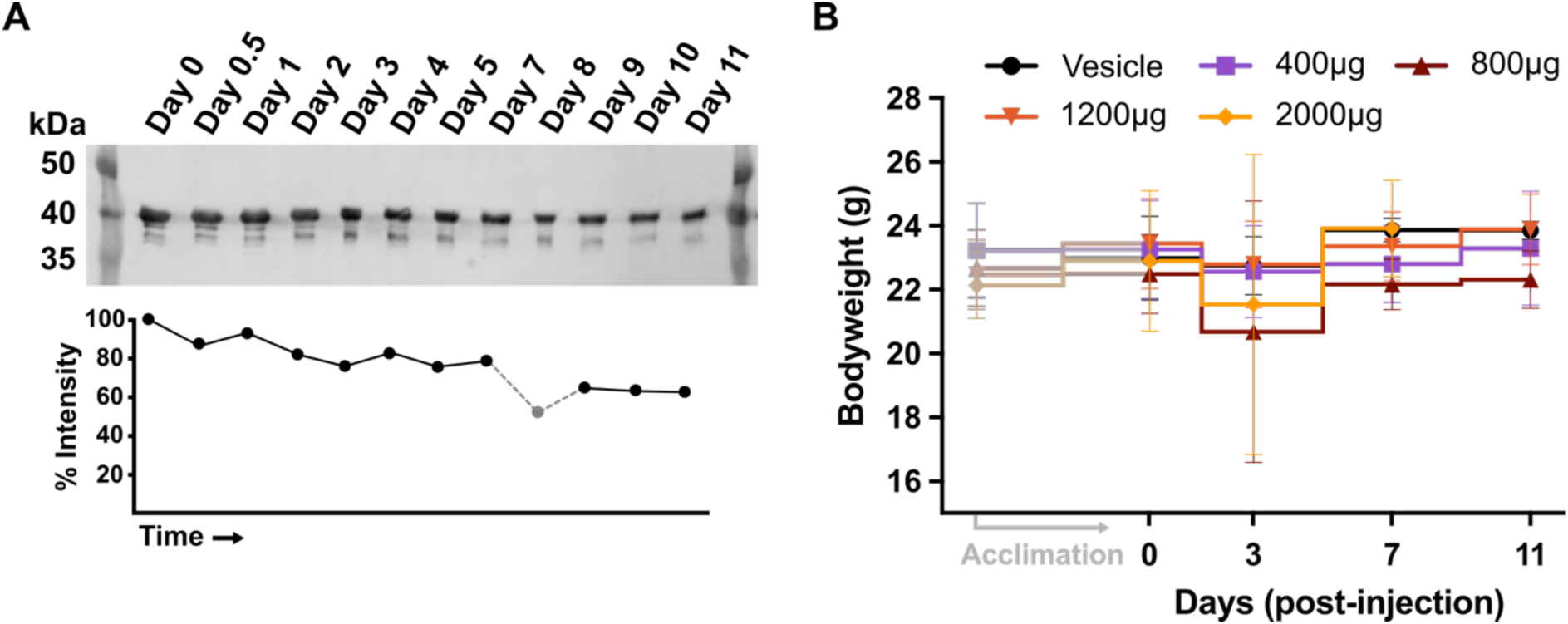
SH-01 has high plasma stability *ex vivo* and shows no acute toxicity in mice. **A** Western blot analysis of *ex vivo* stability assay of SH-01 in human plasma shows that after 11 days, 63% of the protein remains intact. **B** Line plot illustrating total body weight (BW) of C57BL/6 mice immunized with various SH-01 doses, ranging from 400 to 2000 µg per 30g BW. The BW at the beginning of the acclimation period has been included for reference. Over 11 days, no significant changes in BW could be observed across all doses (n = 3; data points are shown as mean values ± SD).

Maximum-tolerated dose studies with SH-01 were conducted in C57BL/6 mice. Endotoxin-free SH-01 was injected intraperitoneally (i.p.) to achieve a plasma concentration of 60 mg/L at the highest dosage, a level sufficient to fully saturate complement factor 3 in the circulatory system. Body weight of the mice was monitored and assessed regularly. None of the dose groups exhibited significant changes in body weight when compared to the control group (Figure 6B). Animals were euthanized on day 12 and gross necropsy revealed no pathological changes to any organ across all dose groups.

### SH-01 prevents hemolysis of PNH erythrocytes similarly to Eculizumab

The absence of negative complement regulators CD55 and CD59 renders the erythrocytes of PNH patients highly susceptible to hemolysis due to the unregulated activity of the AP (Figure 1). Eculizumab, a therapeutic antibody and the current gold standard for treatment of PNH, binds to complement factor C5 at the beginning of the TP, thereby preventing the formation of the MAC and subsequent cell lysis. This, however, comes at the cost of blocking all three complement pathways, leaving patients vulnerable to opportunistic infections that would otherwise be intercepted by the action of the classical and lectin pathways(*35*).

Naturally, we investigated whether SH-01, which selectively inhibits only the AP, can prevent hemolysis of isolated erythrocytes from PNH patients in a manner similar to eculizumab. Erythrocytes for flow cytometry analysis were obtained from blood samples taken from PNH patients undergoing eculizumab treatment (patient A & B, blood was drawn immediately before administration of the next maintenance dose) and untreated patients (patient C).

Fluorescent staining with an anti-CD59-AF_405_ antibody allowed to distinguish between healthy cell populations (CD59^+^) (Figure 7, Q1+Q2) and those affected by the deficiency (CD59^-^) (Figure 7, Q3+Q4). Double-staining with a compatible, anti-C3c-FITC antibody allowed for simultaneous observation of C3b deposition (Figure 7).

**Figure 7.**
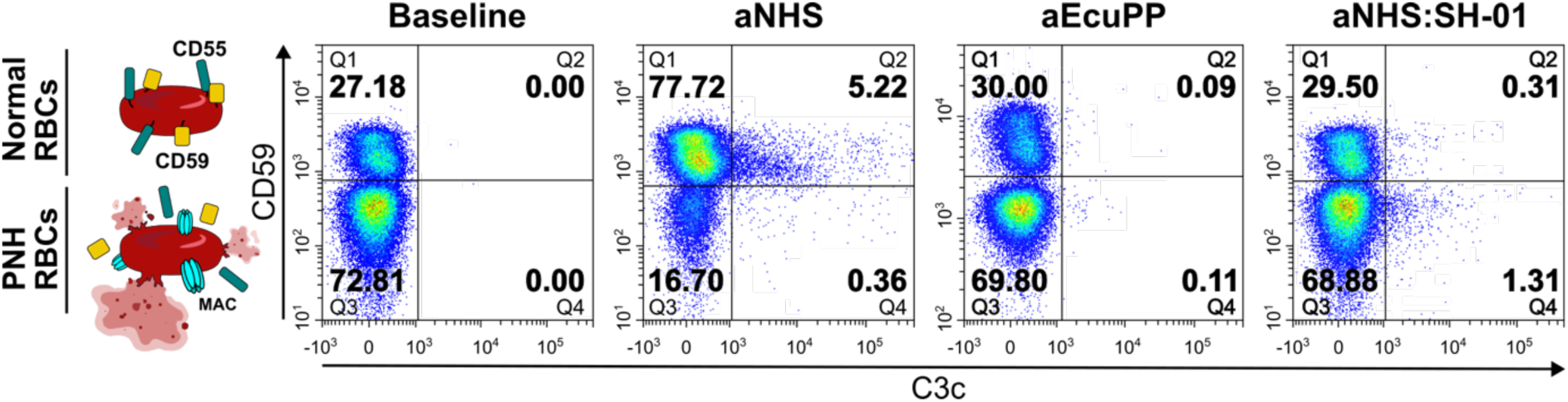
SH-01 effectively prevents lysis of PNH erythrocytes. Flow cytometry analysis of hemolysis and C3b deposition on PNH patient erythrocytes. Two-parameter density plots, divided into four quadrants, illustrate presence of CD59 and C3c on single, intact RBCs. Normal cells (CD59^+^) are shown in Q1+Q2, PNH phenotype cells (CD59^-^) in Q3+Q4. C3b-positive cells are found in Q2+Q4. The percentages of RBCs in each quadrant are indicated. Incubation in acidified human serum (aNHS, triggering AP activation) leads to lysis of most CD59^-^ cells. Hemolysis of erythrocytes incubated in presence of SH-01 and eculizumab is effectively reduced to baseline levels.

Baseline levels of red blood cells (RBCs) with CD59^-^ phenotype were established by incubation of erythrocytes in PBS. For Patient A, flow cytometry analysis revealed 72.81% CD59⁻ cells and 27.18% cells expressing CD59⁺ on their surface. In the absence of human serum, both populations were negative for C3b after washing.

Incubating patient erythrocytes with acidified, ABO-matched human donor serum (aNHS) resulted in rapid lysis of CD59⁻ erythrocytes, as shown by reduction to only 17.06% (16.7% C3c^-^ + 0.36% C3c^+^) of this cell type. CD59⁺ cells were not affected by cell lysis and constituted the majority of intact cells with 82.94% (77.72% C3c^-^ + 5.22% C3c^+^) (Figure 7). Even though 5.22% of ‘healthy’ cells were positive for C3b deposition, the presence of negative regulators such as CD59, which inhibits formation of the MAC, prevents hemolysis (Figure 7).

As expected, PNH erythrocytes incubated with patient serum containing eculizumab showed no significant cell lysis. Comparable to cells incubated in PBS, 69.91% (69.80% C3c^-^ + 0.11% C3c^+^) of intact RBCs were CD59⁻ cells (Figure 7). Analogously, adding SH-01 to donor-matched serum efficiently prevented hemolysis of CD59⁻ erythrocytes (Figure 7). Flow cytometry analysis revealed a fraction of 70.19% CD59⁻ (68.88% C3c^-^ + 1.31% C3c^+^) cells, indicating that these cells were not vulnerable to cell lysis in the presence of SH-01. For both CD59⁺ and CD59⁻ cells, small populations (0.31% and 1.31%) were identified as positive for C3b. In these cases, SH-01 did not fully prevent C3b deposition but likely prevented lysis due to its ability to block AP C5 convertase activity. However, it should be noted that in presence of SH-01, only 1.62% of all intact cells (0.31% CD59^+^ + 1.31% CD59^-^) were positive for C3b deposition, a 3.4-fold reduction from 5.58% (5.22% CD59^+^ + 0.36% CD59^-^) of cells incubated in aNHS. This reduction is likely underestimated because most intact cells observed after aNHS incubation are healthy erythrocytes (CD59+), which naturally degrade surface-bound convertases through CD55 and prevent MAC assembly via CD59. Furthermore, in the absence of SH-01, C3b is rapidly converted to iC3b and subsequently degraded to C3d,g, which is not detected by the anti-C3c-FITC antibody. Therefore, the inhibitory effect of SH-01 on C3b deposition is not accurately represented by the apparent 3.4-fold reduction from aNHS.

Comparable results were obtained for analyses performed with erythrocytes from patients B and C (Supplementary Figure 6). However, patient C did not undergo eculizumab treatment, resulting in a lower initial number of intact CD59⁻ PNH erythrocytes. This finding also showed that prior treatment with eculizumab was not necessary for SH-01 to efficiently prevent hemolysis (Supplementary Figure 6).

## DISCUSSION

Previously, we described the structure of ISG65 and its ability to inhibit the alternative complement pathway by binding to complement factor C3b(*28*). In this work, we elucidated the extended mechanism of selective AP inhibition by SH-01 further and demonstrated that SH-01 effectively inhibits the lysis of PNH erythrocytes, similar to eculizumab. Furthermore, we show that SH-01 exhibits high plasma stability, no acute toxicity and unique structural properties that minimize its immunogenic footprint. With its distinctive mechanism of action, SH-01 represents a new class of C3-based alternative pathway inhibitors, potentially useful not only for the treatment of PNH, but also for other complement diseases characterized by a dysregulated alternative pathway.

While we could previously demonstrate that SH-01 selectively inhibits the AP, but not the CP/LP, we were unable to propose a complete mechanism as inhibition of the AP C5 convertase only accounts for a partial abrogation of cell lysis via the AP(*28*). In contrast to the CP and LP, the alternative pathway is not triggered by microbial surface patterns but instead is activated by spontaneous hydrolysis of C3 in solution, leading to the formation of fluid-phase C3 convertases. In the absence of local concentration maxima of C3b, generated by surface-bound CP/LP C3 convertases near the membrane, the deposition of C3b generated via the AP can be effectively prevented by SH-01.

How exactly does binding to SH-01 prevent C3b deposition? As previously noted, in addition to the primary interaction interface with the TED of C3b, SH-01 forms an additional, secondary interface between its disordered head domain and the CUB linker of C3b(*28*).

The CUB domain serves as a flexible tether, allowing movement of the TED relative to the MG ring, thereby facilitating proper orientation of Gln1013 of the newly exposed thioester for conjugation to hydroxyl-groups on the cell surface(*36–38*). Simultaneous binding of the CUB and TED via SH-01 could severely restrict the conformational flexibility required to position the conjugation-competent glutamine residue correctly, thus effectively inhibiting deposition of C3b (Supplementary Figure 7). Even if a small amount of C3b is deposited on cell surfaces despite the presence of SH-01, SH-01 can inhibit the progression of the cascade regardless, as previously demonstrated, by inhibiting the AP C5 convertase C3bBbC3b. This secondary mechanism was exemplified in our flow cytometry analysis of erythrocytes treated for paroxysmal nocturnal hemoglobinuria (PNH) (Figure 7).

In addition to our previously incomplete understanding of the precise inhibitory mechanism, the fate of C3b generated by the CP/LP C3 convertase C4bC2a in the presence of SH-01 had remained unexplored. Upon proteolytic cleavage through the membrane-associated convertases, newly generated C3b is already present in close proximity to the cell membrane, likely accumulating at a relatively high local concentration. We postulate that this combined effect could facilitate rapid binding of C3b to the erythrocyte membrane, effectively escaping engagement of fluid-phase SH-01, and thus resulting in the observed deposition following the activation of the CP (Figure 2).

Furthermore, we could show here that SH-01 prevents degradation of C3b to iC3b, most likely through sterically blocking fH binding sites on the TED. Consequently, the presence of SH-01 not only preserves the lytic activity of the CP/LP, but also maintains functionality of the amplification loop when the complement cascade is initiated via the CP or LP (Figure 1).

Its unparalleled specificity for the affected pathway and unique two-stage mechanism set SH-01 apart from eculizumab and other alternative pathway inhibitors, such as iptacopan (Fabhalta, Novartis) and pegcetacoplan (Empaveli, Apellis Pharmaceuticals), both of which have recently received approval by the FDA (U.S. Food and Drug Administration) for the treatment of PNH. Iptacopan, a factor B inhibitor, and pegcetacoplan, a C3 inhibitor, not only stall initiation of AP, but also block the amplification loop after activation of the CP/LP through pathogen recognition, resulting in a weakened innate immune response (Figure 8). Likewise, the C5 inhibitor eculizumab restricts progression of the terminal pathway, thus abrogating complement activity entirely, leaving patients with increased susceptibility to bacterial infection (Figure 8).

**Figure 8.**
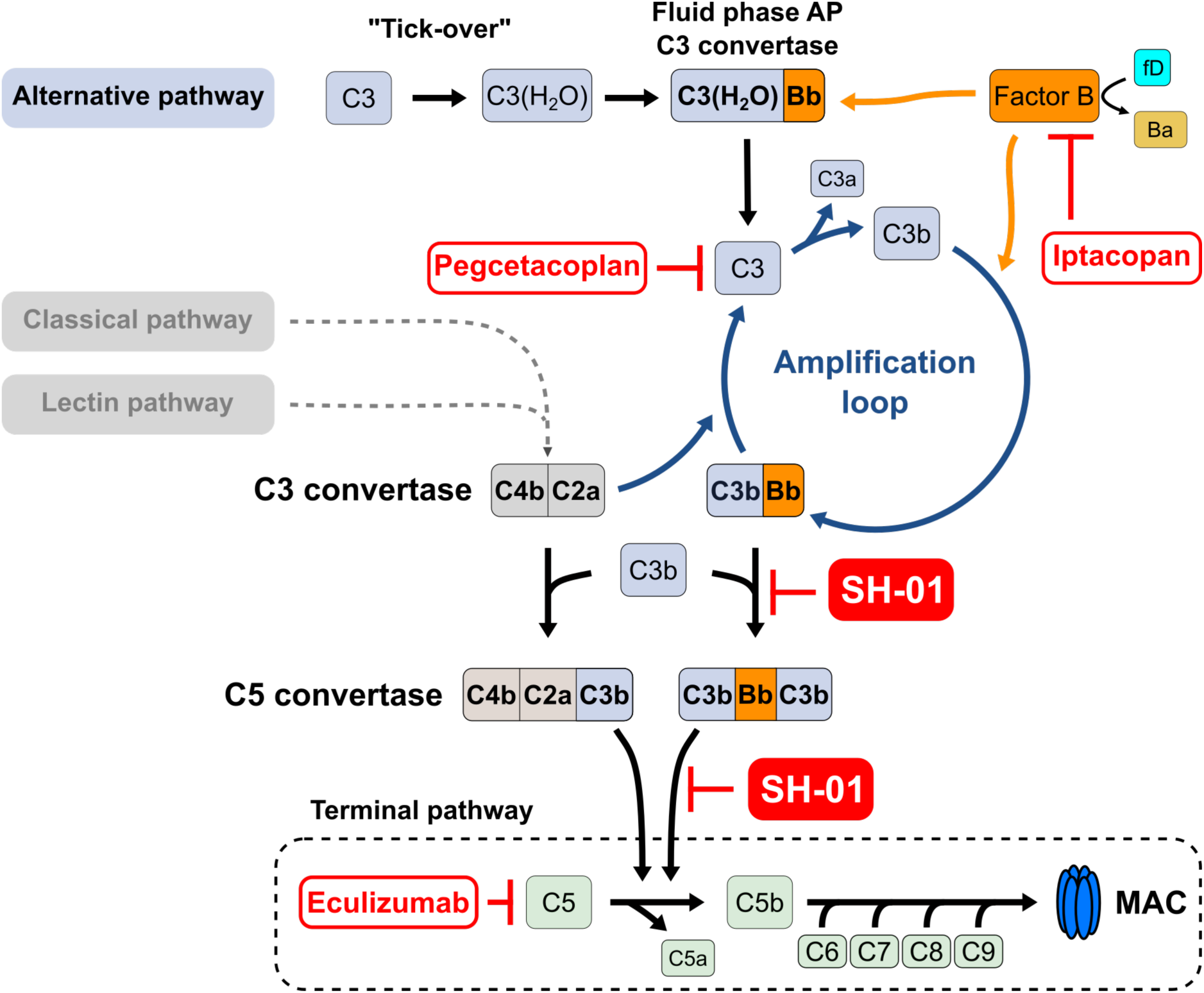
Treatment of alternative pathway dysregulation by complement inhibitors. Iptacopan, an inhibitor of factor B which prevents formation of the mature AP C3 convertase, and pegcetacoplan, an inhibitor preventing proteolytic cleavage of C3 to C3b, not only stall initiation of the AP, but also block the amplification loop after activation of the CP/LP. Eculizumab inhibits C5, the first component of the terminal pathway and point of convergence of all three complement pathways. SH-01 specifically targets the AP by inhibiting the deposition of C3b (not depicted here) and blocking activity of the AP C5 convertase. SH-01 does not impair activation, progression or amplification of the CP and LP.

While iptacopan and pegcetacoplan impede the progression of the AP by inhibiting the proteolysis of crucial components already at its inception, SH-01 does not interfere with the cleavage of native C3 and thus allows generation of C3b and C3a cleavage products (Figure 8). Instead, it prevents the surface deposition of C3b directly, which may be of particular interest for the treatment of complement diseases such as C3 glomerulopathy (C3G), where surface deposition of C3b rather than unregulated cell lysis is responsible for disease manifestation(*35*).

Furthermore, the anaphylatoxins C3a and C5a (generation of both C3a and C5a is abrogated by eculizumab, pegcetacoplan and iptacopan, eculizumab only prevents generation of C5a) are important immunomodulatory protein fragments with a diverse set of functions in the inflammatory response(*39*). While insight into their importance is only emerging, the usage of minimally invasive complement inhibitors with the least possible impact on the overall function of the immune system appears desirable.

Biologics of non-human origin can cause adverse immune reactions upon administration, leading to the rapid neutralization of the drug in the patient’s bloodstream. Due to its origin from the surface of a blood-dwelling human parasite, SH-01 exhibits certain evolutionary traits that enable it to evade immune detection while still exerting its function. In this study, we aimed to characterize the immune-evasive properties of SH-01 in order to harness them for therapeutic use.

Given the large interface between SH-01 and C3b, it appeared conceivable that antibody binding to this region of SH-01 might neutralize the interaction. By employing a flow cytometry-based competition assay, we could show that this is not the case and shielding of the TED binding site does not lead to significantly reduced antibody binding (Figure 3). These findings strongly suggest that the majority of the antibody response is not targeting the interaction interface. In contrast, it is tempting to speculate that the head domain of SH-01 functions as a shield preventing further epitope recognition by antibody-presenting B-cells (Figure 3). Extensive, unstructured loop regions, such as the ones constituting the head domain of SH-01, tend to be more immunogenic than their well-structured counterparts due to their good accessibility and conformational flexibility, which enhances the probability of antibody recognition(*40*). Consequently, we hypothesized that the head domain of SH-01 may serve as an immunodominant decoy to deter antibody binding to the C3b interaction interface and that by blocking this particular region, the immunological signature of SH-01 can be diminished.

While antibody binding to the head domain of SH-01 proved non-neutralizing, we did notice that it induces large-scale conformational changes that result in an increased uptake of deuterium in HDX-MS experiments (Figure 5). This allosteric effect, which does not diminish the interaction of SH-01 with the TED of C3b, could have two conceivable consequences. On the one hand, it may result in an expansion of the low-complexity regions beyond the head domain. An increase in immunogenic, non-functional epitopes could serve to further divert the antibody response away from the binding site(*40*). Simultaneously, the conformational flexibility and transient nature of extensive low-complexity regions are likely to increase the number of epitopes that lack the necessary features to induce a robust antibody recognition. Loop structures, in particular, frequently exhibit significant flexibility, enabling them to adopt a wide range of different conformations. When a loop’s structure deviates from the previously recognized conformation, it may result in impaired antibody binding. This phenomenon could explain both the low number of AF594^+^ B-cells in our flow-cytometry analysis and the emergence of only one high-affinity IgG from our hybridoma selection when using SH-01 as an immunogen.

On the other hand, the induction of large-scale conformational changes could eliminate previously targeted epitopes, thereby reducing antibody recognition. This, in conjunction with the emergence of neo-epitopes in SH-01 upon antibody binding to the head domain, may suppress the overall immune recognition of SH-01.

In conclusion, we could demonstrate here that SH-01 does not elicit a neutralizing immune response that disrupts the interaction with C3b and that antibodies predominantly bind to an immunodominant decoy region, thereby allosterically modulating SH-01 and further contributing to the immune evasion of the latter. These findings underline the potential for the therapeutic use of SH-01 and furthermore suggest that Fab 3E12, or any other antibody(fragment) recognizing a similar epitope in the head domain, may be used as a potential supplement to SH-01. When applied in conjunction with the drug or administered as a booster, it could serve to reduce the immunogenicity of SH-01, either by simply concealing immunogenic regions or by introducing neo-epitopes that diminish the adaptive immune response.

While SH-01 does not elicit a neutralizing antibody response and has proven stable in human plasma over prolonged periods of time, biologics smaller than 30-50 kDa are effectively filtered from the circulatory system through renal clearance(*41*). Rapid removal consequently reduces their effective contact time with the target proteins in the bloodstream, necessitating more frequent administration of the inhibitor. With a molecular weight of 38 kDa, SH-01 is likely to be affected by this elimination mechanism. To increase the serum residence time of SH-01, its overall size could be increased by fusion to the Fc part of an antibody or to serum albumin(*42, 43*). These strategies and their successful implementation have been previously described and should be considered for future pharmacokinetic studies of SH-01 during its pre-clinical development.

In recent years, a growing body of evidence indicating the importance of the complement system in the pathogenesis of variety of severe diseases has spurred the development of several novel complement inhibitors. However, due to the diverse nature of disease pathologies and their underlying molecular dysregulation, no single drug can serve as a universal cure.

Moreover, complex diseases with complement contributions, such as preeclampsia, Alzheimer’s disease, and some forms of cancer, will likely require combination therapies(*35, 44*). Such therapies would greatly benefit from therapeutics with different, complementary modes of action, such as SH-01.

## Acknowledgements

We acknowledge CMS-Biocev (“Biophysical Techniques, Crystallization, Diffraction, Structural Mass Spectrometry”) of CIISB, Instruct-CZ Centre, supported by MEYS CR (LM2018127) and CZ.02.1.01/0.0/0.0/18_046/0015974, for MS measurements conducted by Petr Pompach. We thank the beamline staff at the BioSAXS beamline P12 at DESY Hamburg for their assistance during measurements. Our thanks also go to Petr Mlejnek from the Department of Biological Controls, IPHYS CAS, in Prague, Czech Republic, for performing mouse immunizations and toxicology studies, and to JPT Peptide Technologies in Berlin, Germany, for peptide microarray mapping. Research in SZ’s lab was supported by the Czech Science Foundation (project 22-21612S). HS was supported by the Grant Agency of Charles University (project no. 383821/2600).

## Author contributions

HS and SZ conceptualized the study and conducted protein expressions and purifications. SAXS measurements were performed and evaluated by HS and SZ. FACS measurements were conducted and evaluated by HT, with input from MH, MZ and HS. JC provided patient samples, and AK organized blood collections. JV produced hybridoma cell lines with assistance from SZ. PP conducted the HDX-MS experiments. HS and SZ wrote the manuscript, incorporating input from MZ, and later revised and edited it.

## Notes

### Competing Interest Statement

The authors have declared no competing interest.

