## supplementary information for "A selective alternative pathway complement inhibitor for treatment of paroxysmal nocturnal hemoglobinuria"

A

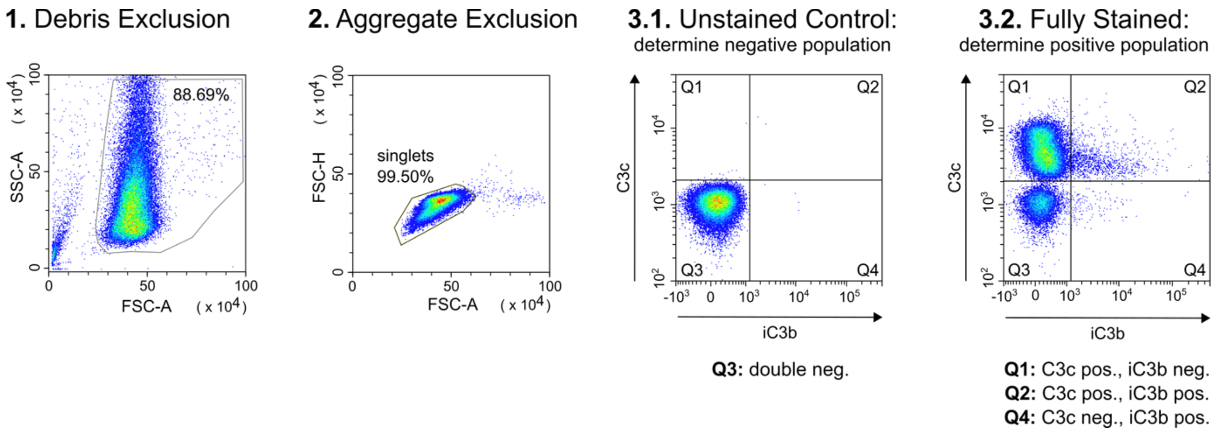

B

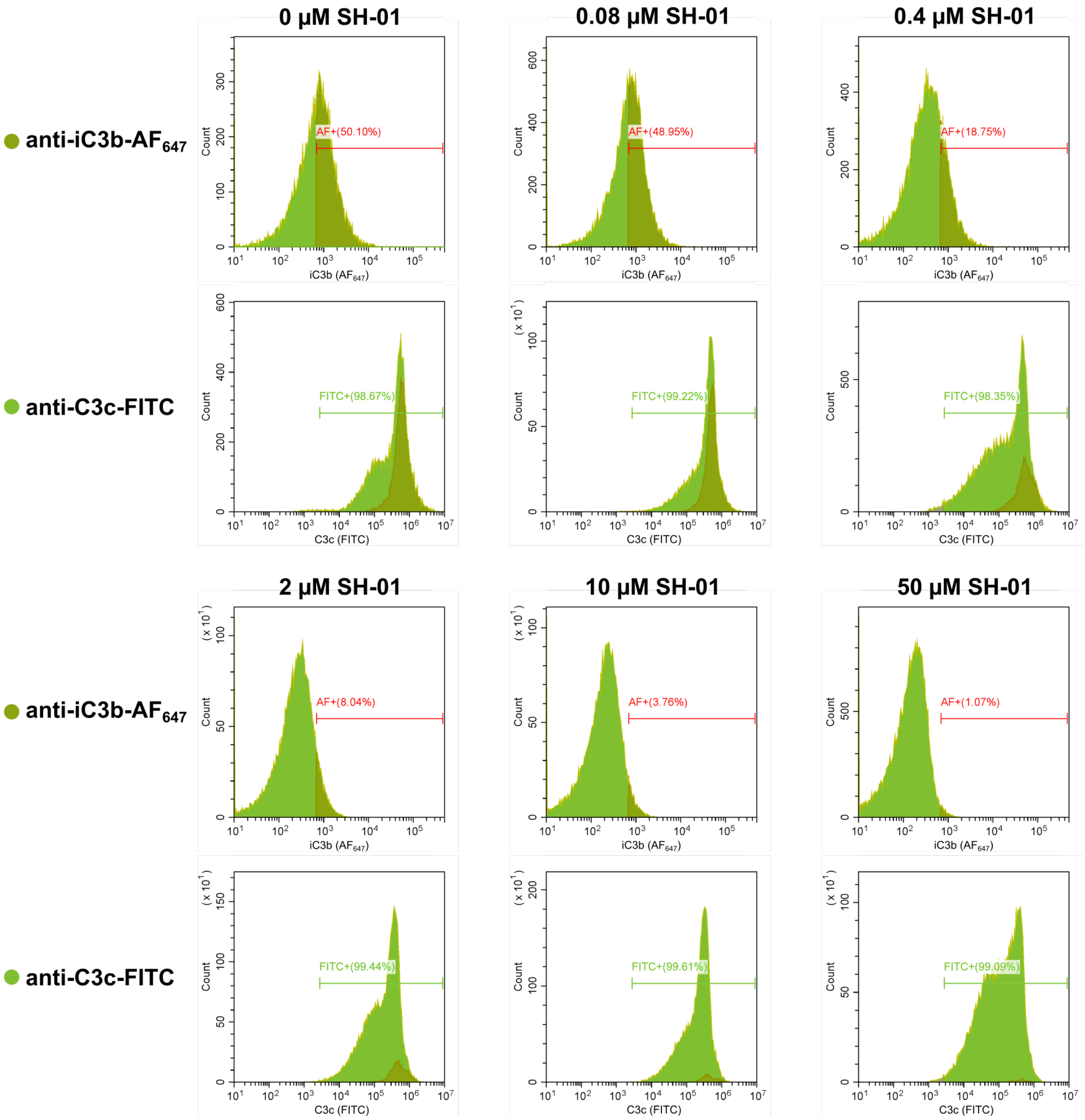


**Supplementary Figure 1. Flow cytometry analysis of C3b deposition on red blood cells during CP/LP.**

**A** Employed gating strategy. Initially, cell debris was excluded based on a side scatter area (SSC-A) vs. forward scatter area (FSC-A) density plot (1). Next, single cells were gated on a FSC-A vs. forward scatter height (FSC-H) density plot (2). Two-parameter density plots, divided into four quadrants, were subsequently utilised to analyse the presence of C3c and iC3b signal on single cells. C3c was stained with a polyclonal anti-C3c-FITC IgG (ab4212, Abcam) (1:300 final dilution) and detected in the FITC (or green) channel. iC3b was stained with a monoclonal anti-iC3b antibody (A209, Quidel), conjugated with Alexa Fluor® 647 (ab269823, Abcam) (2.4 ng/µL final concentration) and detected in the APC (or red) channel. Unstained RBCs were used to establish the level of background fluorescence and to define the negative cell population (3). Presence of iC3b and C3c was evaluated on fully stained cells, allowing for the identification of single positive, double positive, and double negative cell populations (4). 50 000 events were collected and are depicted as density plots. **B** Flow cytometry histograms for APC and FITC channels at various concentrations of SH-01. Each histogram depicts an overlay of anti-iC3b-$AF_{647}$ (dark green) and anti-C3c-FITC (light green) positive cells. Respective gates with percentage of positive, single cells are indicated with a red and green line. It should be noted that the percentages shown here deviate marginally from values reported in Figure 2 due to small differences in manual gate placement.

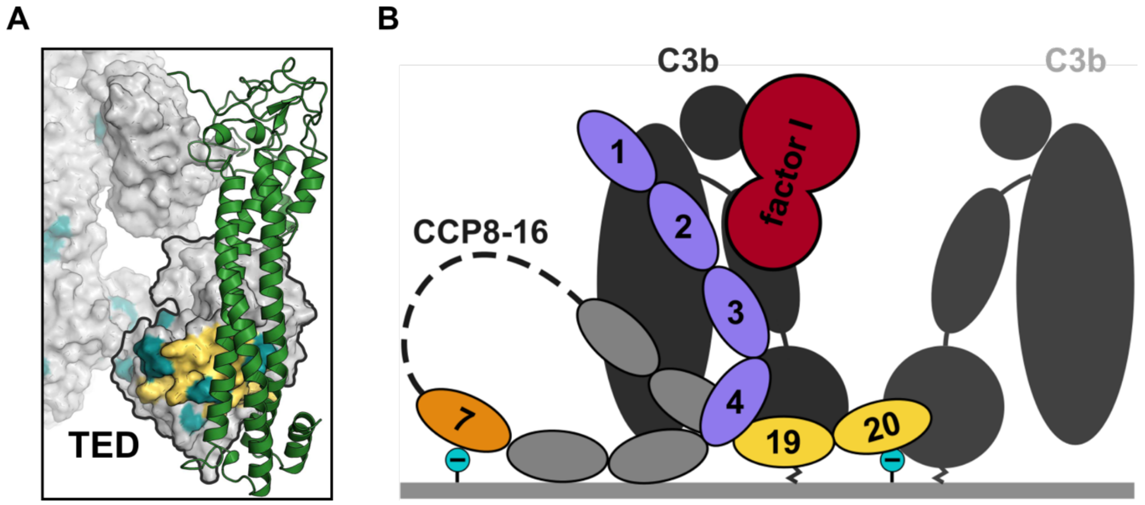


**Supplementary Figure 2. SH-01 occupies binding site of factor H CCP19-20 on the TED of C3b.**

**A** SH-01, depicted in green cartoon illustration, overlaps with the fH binding sites CCP 19-20 on the TED (black outline) of C3b (grey). The fH binding site on C3b is coloured in yellow, while turquoise regions indicate point mutations in C3b, leading to decreased affinity for fH, identified in patients with aHUS disease. **B** Schematic representation of fH and fI assembly on surface-bound C3b. Cell surface attachment via TED is depicted as a serrated line. fH CCPs with explicit roles in C3b binding are highlighted. CCP 19 is a key mediator for correct orientation of fH on surface-bound C3b by binding to the TED. CCP 20 has been shown to bind neighbouring TEDs, and, together with CCP 7, glycosaminoglycans (GAGs) on the cell surface (cyan). CCP 1-4 (purple) harbour the fH co-factor activity for catalytic activity of fI (red). This assembly is based on experimental data (PDBs 2WII and 2XQW), but due to clashing placements of CCP 4 and 19 it may not be entirely representative of the assembly *in vivo*.

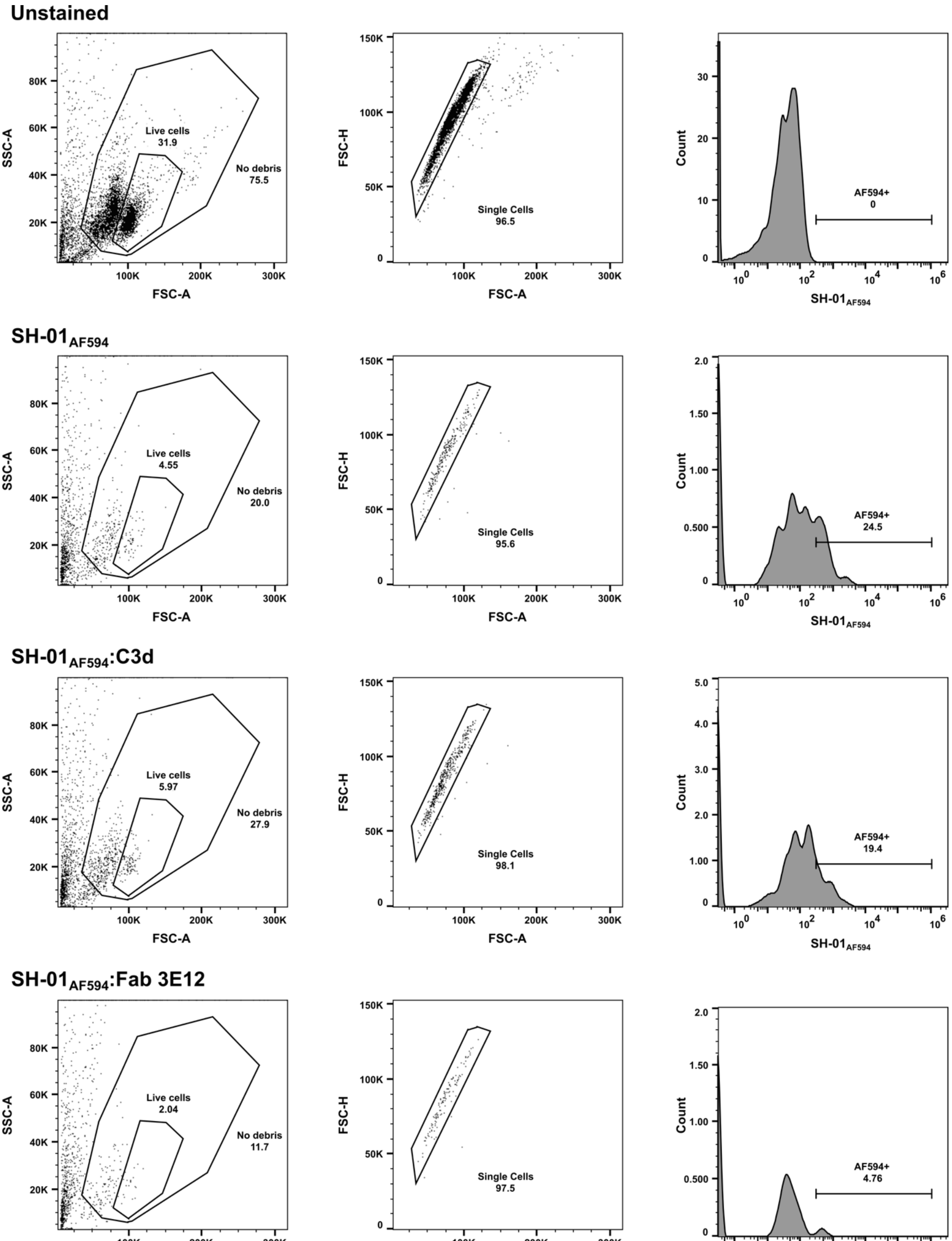


**Supplementary Figure 3. Gating strategy for Flow cytometry-based competition assay.**

$AF_{594}$ positive B-cells were detected using the mCherry channel. Debris was excluded based on SSC-A vs. FSC-A plots, cell viability was assessed using Zombie NIR dye and live cells gated for (dot plot with dead cell exclusion using Zombie NIR not shown, live cell population indicated in SSC-A vs. FSC-A plot). Doublets (aggregates) were discriminated using a FSC-H vs. FSC-A bivariate plot. Incubation of B-cells with PBS buffer instead of SH-01$_{AF594}$ established a fluorescence threshold for $AF_{594}$ positive cells. Addition of SH-01$_{AF594}$ alone resulted in 24.5% of $AF_{594}$ positive B-cells and addition of SH-01:C3d caused no significant decrease in AF594 positive cells. However, addition of SH-01 saturated with Fab 3E12 caused a large decrease in AF594 positive cells to only 4.76%.

**Supplementary Table 1:** Reporting summary for SAS data acquisition, sample details, data analysis, modelling fitting and software used.

| | |
|---|---|
| **(1) Sample details** | |
| Sample | SH-01:Fab 3E12 |
| Organism | *Trypansoma brucei gambiense: Mus muculus* |
| Source | *E. coli* T7 Shuffle : Hybridoma cell line |
| Molecular mass *M* from chemical composition (Da) | ~ 91 000 |
| For SEC-SAS, loading volume/concentration, (mg ml$^{-1}$) injection volume (μl), flow rate (ml min$^{-1}$) | 7.2 mg mL$^{-1}$, 50 μL, 0.3 mL min$^{-1}$ |
| Solvent composition | 20 mM HEPES, 150 mM NaCl, 3% (v/v) glycerol, pH 7.5 |
| **(2) SAS data collection parameters** | |
| Source, instrument and description or reference | BioSAXS beamline P12 at DESY (Hamburg, Germany) with Pilatus6M detector in vacuum |
| Wavelenght (Å) | 1.24 |
| Beam geometry (size, sample-to-detector distance) | 200 x 120 μm , 3.0 m |
| *q*-measurement range (Å$^{-1}$) | 0.23 – 44.3 |
| Absolute scaling method | n/a (no absolute determination of I(0)) |
| Basis for normalization to constant counts | The data were normalized to the intensity of the transmitted beam and radially averaged |
| Exposure time, number of exposures | 1800 successive 0.5 second frames of SEC elution |
| Sample configuration including path length | Quartz glass capillary, 1 mm diameter |
| SEC column type | Superose 6 5/150 GL |
| and flow rate where relevant | 0.3 mL min$^{-1}$ |
| Sample temperature (ºC) | 20 |
| **(3) Software employed for SAS data reduction, analysis and interpretation** | |
| SAS data reduction: | Primus and Chromxis from ATSAS 3.2.1 |
| Calculation of ε from sequence: | ProtParam |
| Calculation of $\Delta\overline{\rho}$ and $\overline{\upsilon}$ values from chemical composition | Primus from ATSAS 3.2.1 |
| Basic analyses: Guinier, *P*(*r*), $V_P$ | Primus from ATSAS 3.2.1 |
| Shape/bead modelling: | DAMMIF amd DAMAVER via ATSAS online |
| **(4) Structural parameters** | |
| Guinier Analysis | |
| *I*(0) (arbitrary) | 49.81 +/- 0.05 |
| $R_g$ (Å) | 38.4 +/- 0.1 |
| *q*-range (Å$^{-1}$) | 0.83 – 3.4 |
| Quality-of-fit parameter (r$^2$) | 0.9986 |
| *P*(*r*) analysis | |
| *I*(0) (arbitrary) | 50 +/- 0.54 |
| $R_g$ (Å) | 40 +/- 0.06 |
| $d_{max}$ (Å) | 140 |
| *q*-range (Å$^{-1}$) | 0.83 – 44.3 |
| *M* from Bayesian inference (Da) | 78500 |
| Porod Volume (Å$^3$) | 123649 |

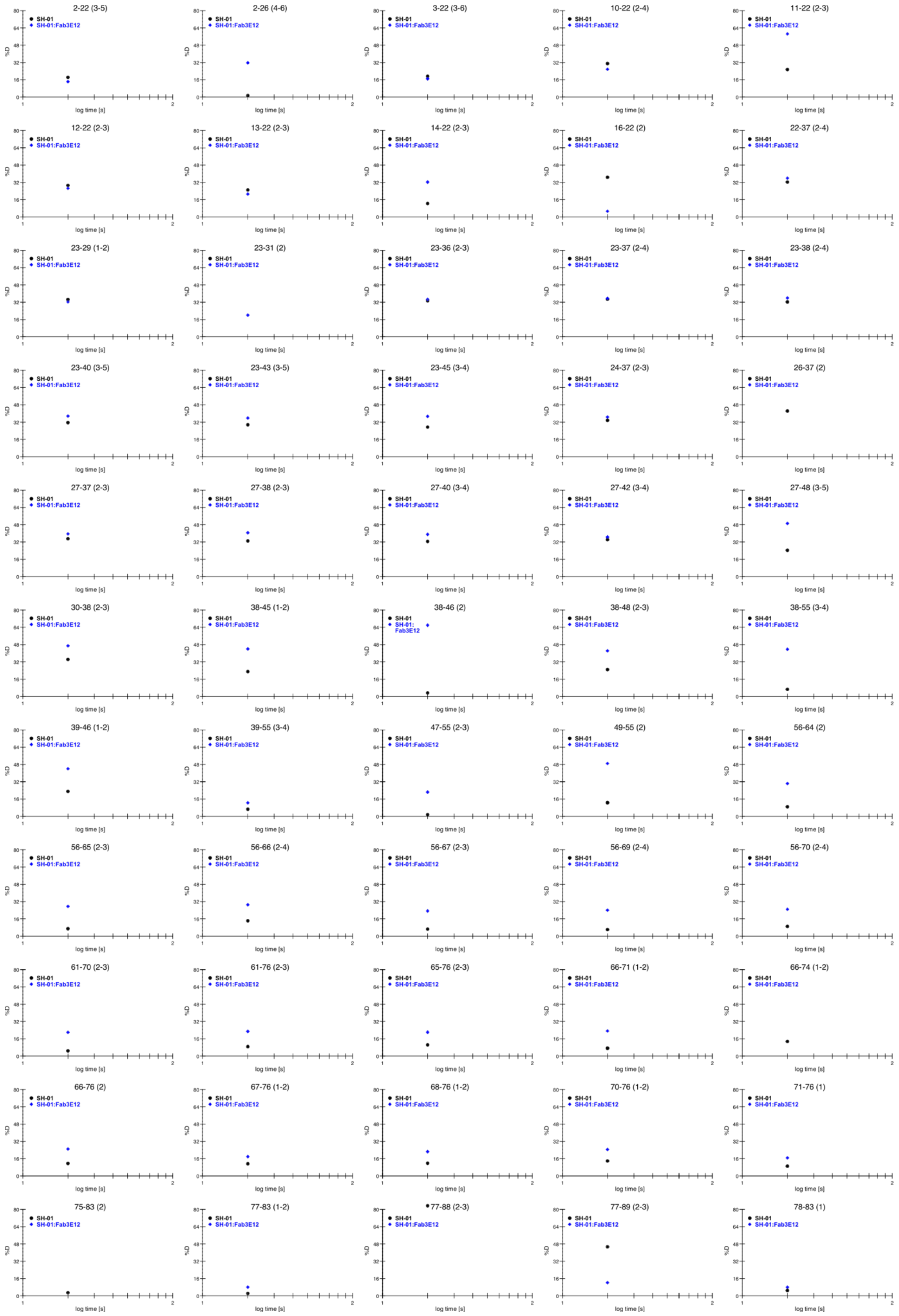
2-22 (3-5)
2-26 (4-6)
3-22 (3-6)
10-22 (2-4)
11-22 (2-3)
12-22 (2-3)
13-22 (2-3)
14-22 (2-3)
16-22 (2)
22-37 (2-4)
23-29 (1-2)
23-31 (2)
23-36 (2-3)
23-37 (2-4)
23-38 (2-4)
23-40 (3-5)
23-43 (3-5)
23-45 (3-4)
24-37 (2-3)
26-37 (2)
27-37 (2-3)
27-38 (2-3)
27-40 (3-4)
27-42 (3-4)
27-48 (3-5)
30-38 (2-3)
38-45 (1-2)
38-46 (2)
38-48 (2-3)
38-55 (3-4)
39-46 (1-2)
39-55 (3-4)
47-55 (2-3)
49-55 (2)
56-64 (2)
56-65 (2-3)
56-66 (2-4)
56-67 (2-3)
56-69 (2-4)
56-70 (2-4)
61-70 (2-3)
61-76 (2-3)
65-76 (2-3)
66-71 (1-2)
66-74 (1-2)
66-76 (2)
67-76 (1-2)
68-76 (1-2)
70-76 (1-2)
71-76 (1)
75-83 (2)
77-83 (1-2)
77-88 (2-3)
77-89 (2-3)
78-83 (1)
SH-01
SH-01:Fab3E12
%D
log time [s]

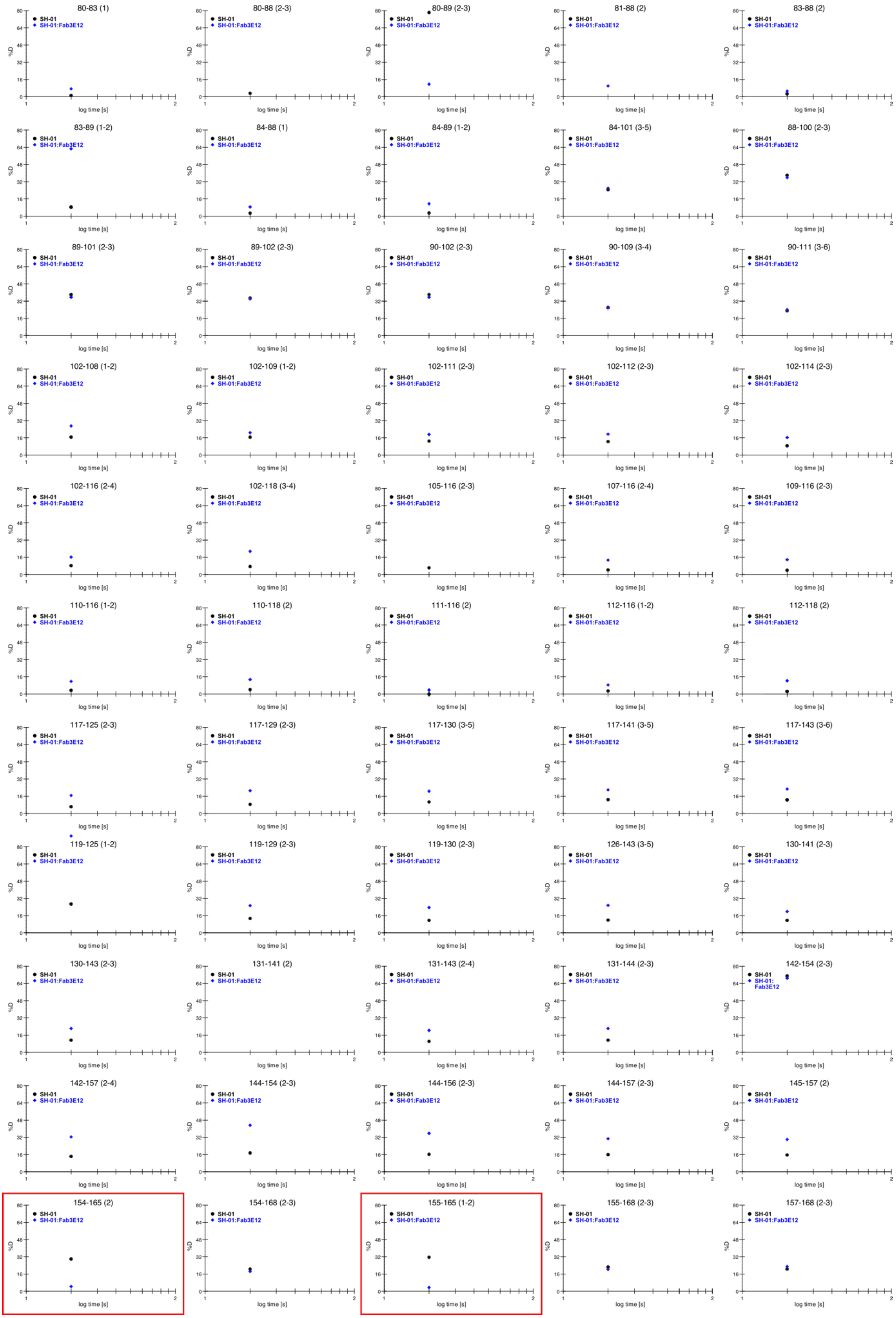
80-83 (1)
80-88 (2-3)
80-89 (2-3)
81-88 (2)
83-88 (2)
83-89 (1-2)
84-88 (1)
84-89 (1-2)
84-101 (3-5)
88-100 (2-3)
89-101 (2-3)
89-102 (2-3)
90-102 (2-3)
90-109 (3-4)
90-111 (3-6)
102-108 (1-2)
102-109 (1-2)
102-111 (2-3)
102-112 (2-3)
102-114 (2-3)
102-116 (2-4)
102-118 (3-4)
105-116 (2-3)
107-116 (2-4)
109-116 (2-3)
110-116 (1-2)
110-118 (2)
111-116 (2)
112-116 (1-2)
112-118 (2)
117-125 (2-3)
117-129 (2-3)
117-130 (3-5)
117-141 (3-5)
117-143 (3-6)
119-125 (1-2)
119-129 (2-3)
119-130 (2-3)
126-143 (3-5)
130-141 (2-3)
130-143 (2-3)
131-141 (2)
131-143 (2-4)
131-144 (2-3)
142-154 (2-3)
142-157 (2-4)
144-154 (2-3)
144-156 (2-3)
144-157 (2-3)
145-157 (2)
154-165 (2)
154-168 (2-3)
155-165 (1-2)
155-168 (2-3)
157-168 (2-3)
SH-01
SH-01:Fab3E12
%D
log time [s]

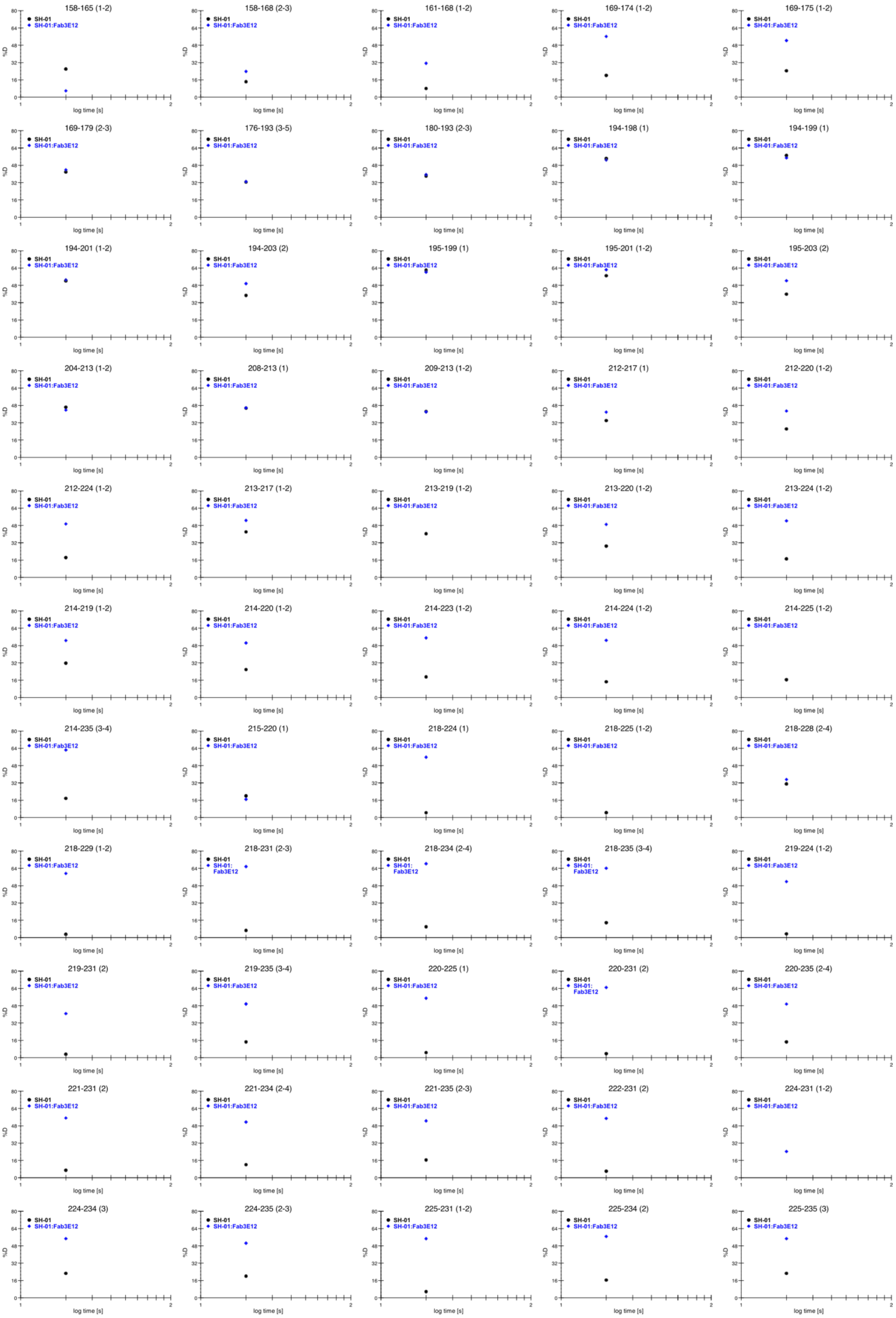
158-165 (1-2)
158-168 (2-3)
161-168 (1-2)
169-174 (1-2)
169-175 (1-2)
169-179 (2-3)
176-193 (3-5)
180-193 (2-3)
194-198 (1)
194-199 (1)
194-201 (1-2)
194-203 (2)
195-199 (1)
195-201 (1-2)
195-203 (2)
204-213 (1-2)
208-213 (1)
209-213 (1-2)
212-217 (1)
212-220 (1-2)
212-224 (1-2)
213-217 (1-2)
213-219 (1-2)
213-220 (1-2)
213-224 (1-2)
214-219 (1-2)
214-220 (1-2)
214-223 (1-2)
214-224 (1-2)
214-225 (1-2)
214-235 (3-4)
215-220 (1)
218-224 (1)
218-225 (1-2)
218-228 (2-4)
218-229 (1-2)
218-231 (2-3)
218-234 (2-4)
218-235 (3-4)
219-224 (1-2)
219-231 (2)
219-235 (3-4)
220-225 (1)
220-231 (2)
220-235 (2-4)
221-231 (2)
221-234 (2-4)
221-235 (2-3)
222-231 (2)
224-231 (1-2)
224-234 (3)
224-235 (2-3)
225-231 (1-2)
225-234 (2)
225-235 (3)
SH-01
SH-01:Fab3E12
%D
log time [s]

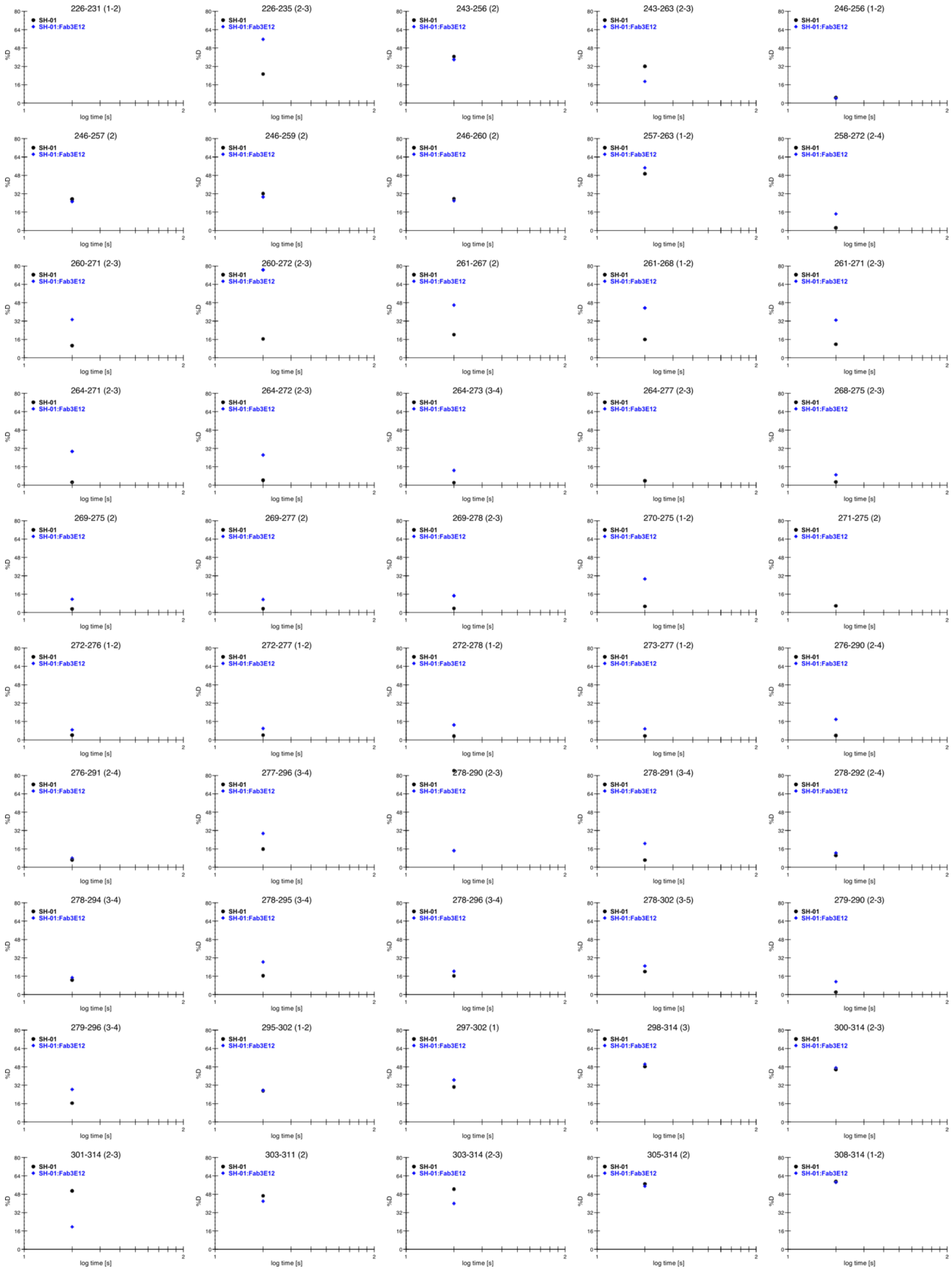


**Supplementary Figure 4. Deuterium uptake plots from HDX-MS analysis of SH-01 in complex with Fab 3E12.**

Deuteration rates of SH-01 (black) and SH-01:Fab 3E12 (blue) after 20s of deuteration are shown. Residue-numbers corresponding to the detected peptide are indicated above the respective plot. Plots corresponding to the TAKSTGL sequence identified as the Fab 3E12 binding site are highlighted with red boxes. Deuteration rates of these peptides was significantly lower in the SH-01:Fab 3E12 complex than for SH-01 alone.

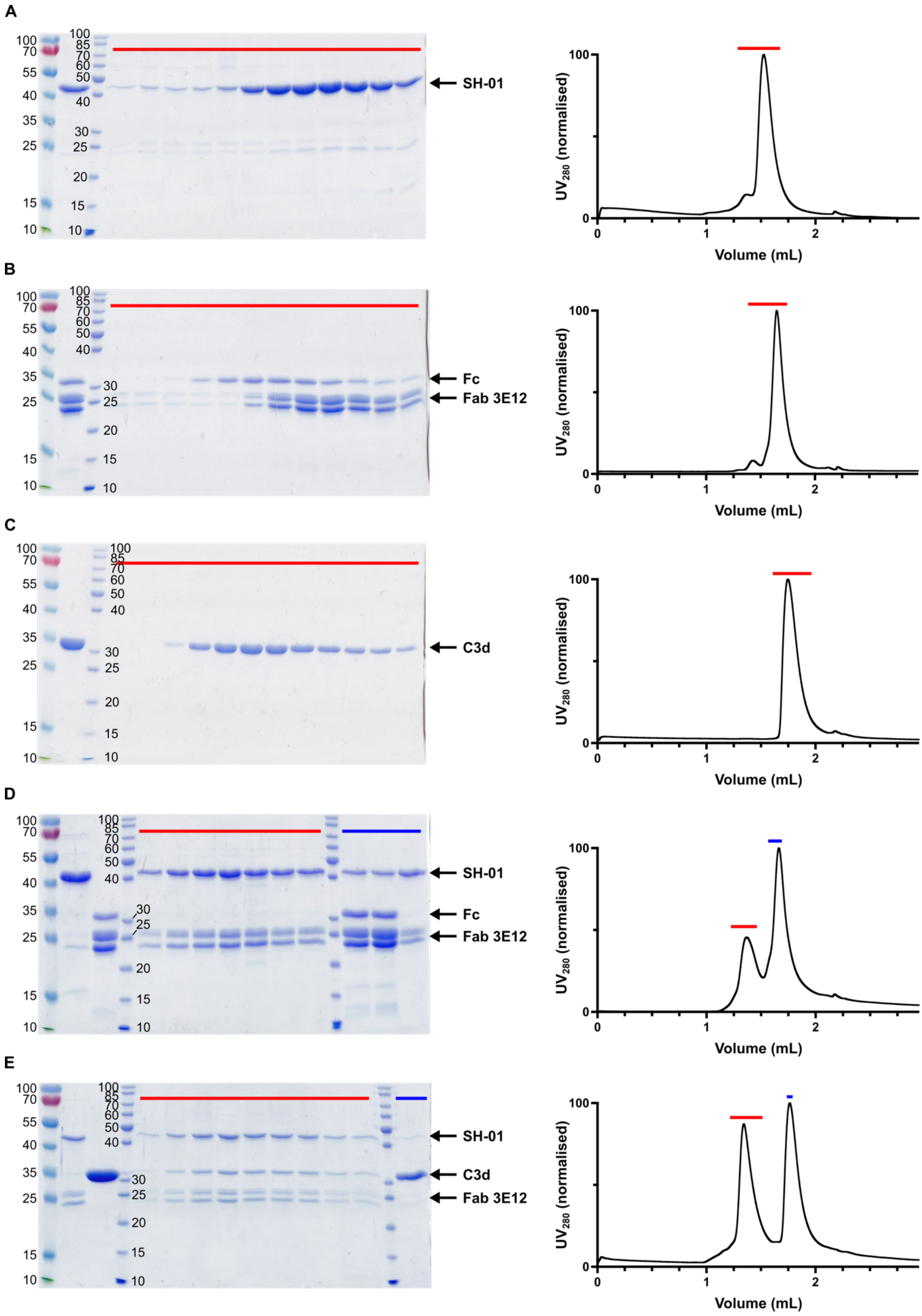


**Supplementary Figure 5. Fab 3E12 binding to SH-01 does not impair its ability to interact with C3d.**
Reducing SDS-PAGE analysis of size exclusion peak fractions and corresponding chromatograms are depicted for individual proteins and their complexes. $UV_{280}$ was normalized to the maximum recorded absorbance for each chromatogram for illustrative purposes. The unedited source data are provided in the Source Data file. Peak fractions visualized in the reducing SDS-PAGE analysis are highlighted in the corresponding section of the chromatograms with red and blue horizontal lines.

Sample lanes on the SDS-polyacrylamide gel without highlights show loading controls of the individual components. The identity of the visualized protein bands is indicated with arrows on each gel. (A) SH-01. (B) Fab 3E12. (C) C3d. (D) SH-01 with a threefold molar excess of Fab 3E12. (E) SH-01:Fab 3E12 complex with threefold molar excess of C3d.

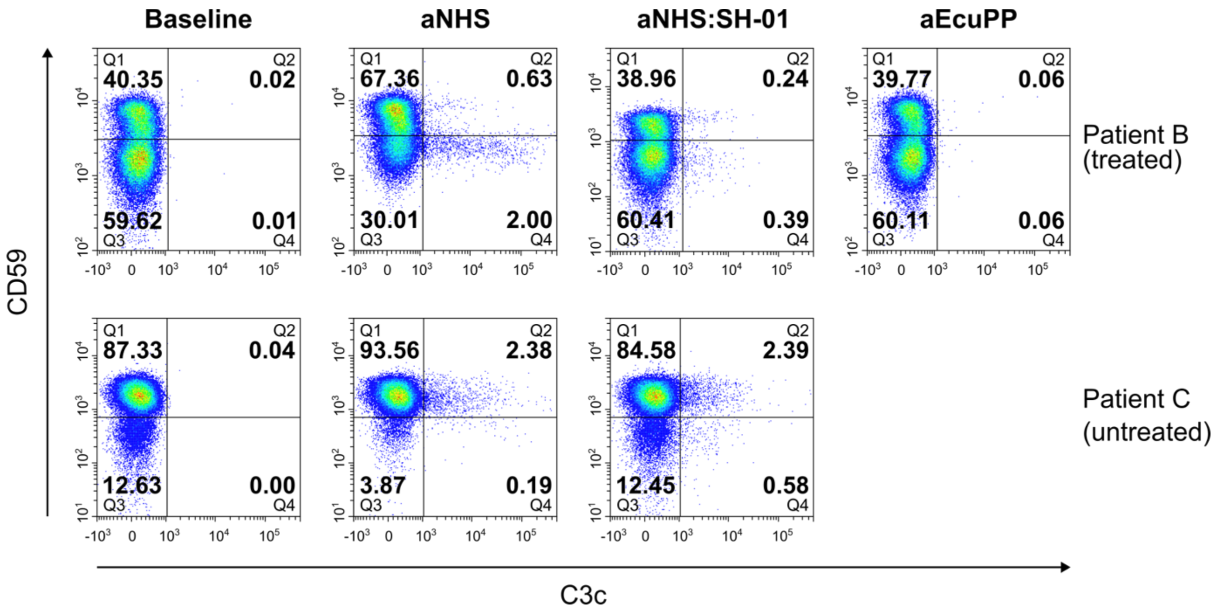


**Supplementary Figure 6. Effect of SH-01 on hemolysis in treated and untreated PNH patients.**

Flow cytometry analysis of hemolysis and C3b deposition on erythrocytes of PNH patients B (treated with eculizumab) and C (untreated). Two-parameter density plots, divided into four quadrants, illustrate presence of CD59 and C3c on single, intact RBCs. The percentage of cells in each quadrant are shown in bold numbering. Baselines and gates were established as depicted in Supplementary Figure 1A but instead of staining for iC3b, anti-$CD59_{AF405}$ monoclonal antibody was used for detection of CD59 expression. Hemolysis and C3 fragment deposition, resulting from the activation of complement alternative pathway, were induced by incubating RBCs in ABO-matched acidified serum supplemented with $MgCl_2$ (aNHS). RBCs from patient C were incubated in NHS pooled from two different donors to mitigate variations in complement activity following the acidification. For patients A or B, NHS from a single donor was used due to the lack of suitable ABO-matched donors. Hemolysis is indicated by a reduction in the number of CD59-negative cells in quadrant Q3. SH-01 (48 μM final concentration), if pre-incubated with NHS prior to serum acidification, effectively protected PNH RBCs from hemolysis (aNHS:SH-01). As a positive control for prevention of hemolysis, RBCs from Patient B were also incubated in patient's own acidified eculizumab-containing plasma (aEcuPP). RBCs were incubated at 37°C for 1h before staining with antibodies. The density plots depict 50 000 events.

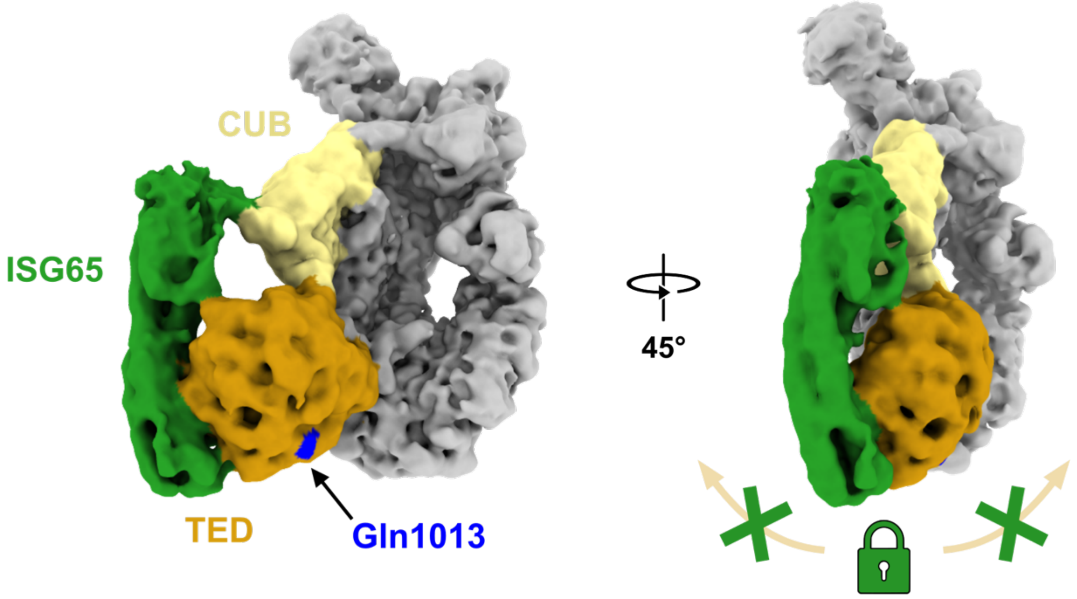


**Supplementary Figure 7. Single-particle cryo-EM reconstruction of ISG65 in complex with C3b.**
Unsharpened cryo-EM density map of an ISG65:C3b reconstruction (EMD-14708). ISG65 (green) contacts TED (orange) and CUB (yellow) simultaneously. This structural arrangement could severely restrict the conformational flexibility required to position the reactive glutamine Gln1013 residue of the thioester (blue) correctly.
